# Viral Protease Nsp5 Hijacks La Autoantigen to Orchestrate the Translation to Replication Transition in SARS-CoV-2

**DOI:** 10.64898/2026.08.05.742966

**Authors:** Risabh Sahu, Harsha Raheja, S N Gagan Gaurav, Santu Paul, Shree V Sabari, Pritam Kumar Ghosh, Raju S. Rajmani, Saumitra Das

**Affiliations:** Department of Microbiology and Cell Biology, Indian Institute of Science, Bangalore-560012; National Institute of Biomedical Genomics, Kalyani, West Bengal

**Keywords:** SARS-CoV-2 lifecycle, La protein cleavage, Nsp5 protease, host virus interaction

## Abstract

Viruses genetically encode proteins to regulate their life cycle. To translate this genetic information, most viruses exploit host translation machinery. Host dependency leads to competition between viral RNA and host mRNAs. Most viruses overcome it by inhibiting cap-dependent translation of host mRNAs and shifting its own translation into cap-independent mode. This switch is largely orchestrated by host RNA binding proteins (RBPs) by binding to viral 5’UTRs. Here we demonstrate that human La autoantigen (La protein) specifically binds to GCAC sequence near the initiator AUG within the 5’UTR of SARS-CoV-2 RNA promoting viral non-structural protein synthesis by cap independent translation, while suppressing replication early in infection. Interestingly, around 18-24 h post-infection, the viral protease Nsp5 cleaves La at a conserved LQ site, abrogating its 5′UTR interaction, which correlates with reduced polysome association and increased viral RNA availability for replication. MD simulation and docking study suggests that the cleavage disrupts La dimerization and truncates its C-terminal intrinsically disordered region, reducing RNA-binding affinity and conformational stability required for translation-competent ribonucleoprotein complexes. Results put forward a unique mechanism by which Nsp5-mediated proteolysis of La drives the temporal switch from translation to replication of SARS-CoV-2.

## Introduction

Viruses encode structural and non-structural proteins to execute their life cycle but heavily depend on the host translational machinery to express their genetic information, creating inherent competition between viral RNAs and host mRNAs. To overcome this constraint, many RNA viruses suppress host cap-dependent translation and redirect ribosomes toward cap-independent mechanisms (1). Internal ribosome entry site (IRES) mediated translation represents one of the most widely exploited cap-independent translation strategies among positive stranded RNA viruses (2). Severe Acute Respiratory Syndrome Coronavirus 2 (SARS-CoV-2), an enveloped, positive sense, single stranded RNA virus is a member of the *Betacoronavirus* genus possesses a 30 kilobase (kb) long RNA genome, which is flanked by 5’UTR and 3’UTR. Following entry, the capped 5’UTR initiates translation of the Open Reading Frame ORF1ab, producing a polypeptide which is proteolytically cleaved to produce 16 non-structural proteins (Nsps) from the genomic RNA, that helps initiating viral RNA replication from 3’UTR (3). Among these, non-structural protein 1 (Nsp1) of SARS-CoV-2 suppresses host cap dependent translation and promotes virus translation by cap independent mechanisms(4). Thus, SARS-CoV-2 reshapes the host translational landscape and relies on cap independent translation to ensure efficient viral protein synthesis at early stages of infection. The structured 5’UTR and 3’UTR of SARS-CoV-2 serve as scaffolds for specifically binding host RNA binding proteins (RBPs). Several high-throughput studies have identified various RBPs, which can positively and negatively regulate virus lifecycle and pathogenesis (5, 6). However, the mechanistic role of host RBPs coordinating the switch between translation and replication remains poorly understood.

In this study, we identify host La autoantigen (Lupus La protein) as a master regulator of SARS-CoV-2 lifecycle. Biochemical screening of all RBPs, using HEK293T-ACE2 S10 extract, revealed enhanced binding of a 52 kDa protein band with the 5’UTR of SARS-CoV-2. Mass spectrometric analysis of the protein bands identified it as the human La protein (La autoantigen). La autoantigen is known to play an essential role in maturation of precursor tRNA to mature tRNA and also shown to bind the IRES element of HCV, CVB3 and polio viruses to regulate their lifecycles (7–11). In this study, a series of biochemical assays validated the interaction of the La protein with the 5’UTR of SARS-CoV-2 RNA. We demonstrate that this protein regulates cap independent translation and replication of viral RNA by binding to the GCAC sequence proximal to the initiator AUG.

Here we demonstrate that La protein enhances cap-independent translation of viral RNA and preferentially promotes viral protein synthesis early in the infection, while negatively regulating viral RNA replication. Further, we observed that one of the viral non-structural proteins, Nsp5 proteolytically cleaves La autoantigen. The cleaved La (N-terminal) fails to interact with 5’UTR of SARS-CoV-2 in the late phase of the virus lifecycle resulting in repression in translation and activation of replication. Taken together, the results uncover a novel interplay between a host RBP (La autoantigen) and virus protease (Nsp5) to dynamically regulate its interaction with the 5’UTR and modulate the temporal switch from virus translation to replication.

## Materials and Methods

### Plasmid constructs

The coding sequence of La protein (1-408aa) cloned between Bam HI and EcoR I sites of pRSET-A was obtained from Duke University, Durham, NC (A kind gift by Dr. Jack Keene). The coding sequence of La was cloned between HindIII and EcoRI sites in pcDNA3.1 HisC for further overexpression-based experiments (pcDNA3.1-La). 5’UTR of pCBB 5’UTR-CoV2 (A kind gift by Dr. Milan Surjit, THSTI) was subcloned in the pcDNA3.1 between NheI and HindIII (downstream of T7 promoter) and was used for *in vitro* transcription. Coding sequence of Wuhan Nsp5 (Nsp5 wild-type) and C145A-Nsp5 (Nsp5 dead mutant) was cloned between NotI and EcoRI of pIRESneo (A kind gift by Dr. Anup Majumder, NIBMG, Kalyani). Coding sequence of Wuhan Nsp3 was cloned between BamHI and EcoRI in pCMV-Tag2B (A kind gift by Dr. Shashank Tripathi, IISc, Bangalore). pcDNA5-CoV2-5’UTR-Rluc and pcDNA5/FRT/TO-Fluc-5’UTR-Rluc were obtained from The Weizmann Institute of Science, Israel (a kind gift by Dr. Rivka Dikstein). Sh scrambled and Sh La constructs were used for silencing experiments.

### Cell lines and transfection

HEK-293T cells expressing ACE2 receptor, Huh7, VeroE6 cells were grown in Dulbecco’s Modified Eagle Medium (DMEM-Sigma) containing 10% Fetal Bovine Serum (FBS, Gibco). Mycoplasma was not detected in the above-mentioned cell lines. For transfection, 1.4 ×10**^5^** HEK293T-ACE2 cells were seeded in each well of 24 well plate. 18 hours of post seeding, transfection experiments were performed in OPTI-MEM (reduced serum medium, Gibco). 200ng to 300ng DNA construct per well of 24 well plate was used for transfection. Lipofectamine-2000 (Invitrogen) was used as a transfection vehicle for transfection experiments as per the manufacturer’s protocol.

### Virus stock preparation and infection

Virus stocks were procured from BEI resources, NIAID, NIH and prepared at Viral BSL3, IISc. Wild-type (WT) virus: Isolate Hong Kong/VM20001061/2020, NR-52282; Delta variant (Lineage B.1.617.2): Isolate hCoV-19/USA/PHC658/2021, NR-55611; Omicron variant (Lineage BA.2.12.1): Isolate hCoV-19/USA/NY-MSHSPSP-PV56475/2022.

Viruses were propagated in Vero E6 cells. Vero E6 cells were infected with WT, Delta and Omicron variants with very low Multiplicity of Infection (MOI=0.01) and 48 hours post infection (h.p.i) supernatants were collected and preserved at −80 °C for further experiments. To titrate the virus, Vero E6 cells were further infected with virus by serial dilution and plaque forming units (pfu) were calculated to determine virus titre by plaque forming assay. HEK-293T-ACE2 cells were infected with virus at MOI=1 for other experiments.

### Dual luciferase assay

Luciferase assays were performed using dual luciferase kit (Promega). To collect Firefly luciferase (Fluc) value, LAR reagent was used and later stop & glow was added to collect Renilla luciferase (Rluc) value using Glomax luminometer (Promega). Fluc construct was used as a transfection control for all luciferase experiments, whereas Rluc was present in the experimental construct. For bicistronic assays, a single construct containing Fluc and Rluc was used and the ratio of Rluc and Fluc reflected cap independent translation of virus.

### Protein purification

*E. coil* BL21 cells were transformed with pRSET-A-La. The protein was purified as described earlier (12). Single colony from the transformed plate was inoculated in 200 mL LB broth containing ampicillin and grown at 37 °C at 180 rpm shaker incubator until OD_660_ reached 0.6. The cultures were induced with 0.6 mM IPTG (isopropyl-1-thio-β-D-galactopyranoside) and placed back in the shaker incubator for 4 hours (37°C). Later, the cells were pelleted and lysed in 5mL lysis buffer (50 mM NaH_2_PO_4_, 300 mM NaCl, 10 mM imidazole, 0.1mM phenylmethylsulfonyl fluoride (PMSF) and 1X bacterial protease inhibitor cocktail, Sigma) by sonication. The supernatant was further incubated with Nickel-NTA (Ni-NTA) agarose slurry and kept for rocking for 4 hours at 4°C. Later, beads were further washed using 200 mL wash buffer (50 mM NaH_2_PO_4_, 300 mM NaCl, 40 mM imidazole) and eluted using elution buffer containing 500mM imidazole. The eluted protein was further dialyzed using dialysis buffer (25 mM Tris, pH 7.4, 100 mM KCl, 7 mM β-mercaptoethanol [β-ME], 10% glycerol) and stored at −80°C.

### *In vitro* transcription (IVT)

pcDNA3.1-5’UTR-CoV2 was linearized using HindIII followed by precipitation using phenol-chloroform method. The linearized DNA was used as a template for *in vitro* transcription. IVT was performed as per the manufacturer’s protocol (Promega, T7 polymerase IVT kit) using ATP, GTP, CTP, UTP (**^32^**P**-**UTP was used for radio-labelled RNA preparation). Reaction mixture was kept at 37°C for 1.5-2 hours following which the RNA was precipitated using 4M ammonium acetate, glycogen and 100% ethanol, and dissolved in nuclease-free water. 1 microlitre of radio-labelled RNA was spotted on DE81 filter paper and washed with phosphate buffer and 100% ethanol and radioactivity was measured using scintillation counter.

### RNA and protein crosslinking by Ultraviolet (UV) irradiation

Radio-labelled 5’UTR-CoV2 RNA and La protein were incubated at 30°C for 30 min in binding buffer (pH 7.6, 25 mM KCl, 3.8% glycerol, 2 mM MgCl_2_, 2 mM dithiothreitol [DTT], and 0.1 mM EDTA, 5 mM HEPES, 1.25 mM ATP, 2Mm GTP). 30 min post incubation, the sample mixture was kept on ice and treated with UV radiation at 254 nm for 20 minutes. Post UV treatment, the mixture was further treated with 50µg of RnaseA (Sigma) for 45 minutes to 1 hour to digest unbound RNA. 1-hour post treatment the samples were heated at 100°C with protein loading dye. The MCS (multiple cloning site) of pGEX vector was transcribed to use as Non-specific RNA (Nsp RNA).

### Surface Plasmon Resonance (SPR)

To perform SPR, SARS-CoV2 5’UTR RNA was biotin labelled by *in vitro* transcription using biotin labelled UTP. Biotin labelled 5’UTR RNA was immobilized on sensor chips (streptavidin coated) to a final concentration of 300 Resonance Units (RU)/flow cell. Interaction between RNA–protein was carried out in a continuous flow of Tris buffer (25 mM Tris (pH 7.5), 100 mM KCl, 7 mM β-mercaptoethanol, and 10% glycerol) at a flow rate of 10 µl/min at 25 °C. Increasing concentrations of 8, 16, 32 and 80 nM of La protein was added on the biosensor chip for 100 seconds (known as association phase) and further kept with only buffer for 300 seconds (known as dissociation phase). Biosensor chip alone (without biotinylated RNA) served as the negative control to normalize with the non-specific background (noise) data. BIA evaluation software (version 3.0) was used to determine the off rate, *k*_d_ (s^-1^) and the on rate, *k*_a_ (M^-1^ s^-1^). Langmuir binding model was used for analysis. The binding affinity, *K_d_* was determined using the following equation: *K_D_* =*K*_d_/*K*_a_.

### IP-RT (Immunoprecipitation followed by real time PCR)

HEK-293T-ACE2 cells were infected at MOI=1 or transfected with Wild-type 5’UTR (GCAC)/Mutant 5’UTR (ACGC) construct. 24 hours or 48 hours post infection/transfection, cell pellet was collected and lysed with polysome lysis buffer (100 mM KCl, 10 mM HEPES pH 7.0, 5 mM MgCl_2_, 0.5 % NP-40, 1 mM DTT, 100 U/mL RNasin, 100U/mL mPIC) at 4°C cyclomixer. Protein G Sepharose beads (G Bioscience) were previously incubated with IP grade La antibody (NBP1-48802, Novus biologicals) overnight. Antibody-bound beads were further incubated with cell lysate overnight and next day beads were washed three times with polysome lysis buffer and then 20% of the beads were further boiled at 100° C for 10 min and immunoprecipitated protein released in supernatant was run on 12% SDS gel and probed with antibody to confirm pulldown. Remaining 80% of the beads were incubated at 50° C after adding 30µg of proteinase K and 0.1% SDS. 30-45min post incubation TRI reagent (Sigma) was added for RNA isolation and checked for association of virus RNA with La protein. IgG was used as negative control for IP experiments.

### RNA isolation followed by quantitative real-time PCR (qRT-PCR)

Total RNA isolated from HEK-293T-ACE2 cells using TRIZol (TRI) reagent (Sigma) as per manufacturer’s protocol. cDNA and qRT-PCR were performed using CoV2 Fwd: 5’ TGTCGTTGACAGGACACGAG 3’, CoV2 Rev: 5’ TTACCTTTCGGTCACACCCG 3’ Actin Fwd: 5’TCACCCACACTGTGCCCA’3, Actin Rev: TGAGGTAGTCAGTCAGGT. Moloney murine leukemia virus (M-MLV) reverse transcriptase was used for cDNA preparation using 600 ng RNA (concentration taken in nanodrop). After cDNA preparation, 2µl of cDNA samples were further used for qRT-PCR using DyNAmo SYBR green (ThermoScientific) as per manufacturer’s instructions. Comparative threshold (CT) value was further used. ΔCT was calculated using CT value of virus RNA and Actin RNA. ΔΔCT was calculated by normalizing ΔCT value of control sample and experimental sample. Fold change was calculated using the formula 2^(-ΔΔCt)^.

### Western Blotting

Protein was isolated from the cell pellets using 1X RIPA (Radio immunoprecipitation) lysis buffer (5X RIPA composition-0.5M Tris-HCL (pH 7.4),1.5M NaCl, 2.5% Na-deoxycholate, 10% NP-40, 10mM EDTA). Concentration of protein lysates were determined using 1X Bradford reagent (Biorad) using manufacturer’s protocol. Equal concentration of proteins (100 µg) was loaded onto 12% SDS gel and then transferred onto a Nitrocellulose membrane (Biorad) or PVDF membrane (MERK). Samples were incubated with specific antibodies to check abundance of different host and virus proteins. Antibodies used for protein detection were anti-SARS-CoV-2 N protein (40143-MM05, Sino Biological), anti-SARS-CoV-2 Nsp5 protein (GTX-135470), anti-SARS-CoV-2 Nsp3 (CST, 88086), anti-SARS-CoV-2 Nsp1 (GTX-135612), anti-La protein (EPR-6570, Abcam), anti-La protein (NBP1-48802, Novus biologicals). Secondary antibodies (anti-mouse IgG HRP tagged, Sigma or anti-rabbit IgG HRP tagged, Sigma) were used before development using Biorad ECL substrates. Mouse-monoclonal anti-β-actin HRP tagged antibody (A3854, Sigma) was used as a control for equal loading of protein lysates.

### Sucrose gradient preparation for Ultracentrifugation

10% and 50% sucrose solution prepared in 1X gradient buffer, (10X gradient buffer: 200mM Tris HCl, 1.5M KCl and 50mM MgCl**_2_**) poured sequentially in the Ultracentrifuge tube (Beckman). To prepare a continuous gradient, samples were spun at 21 rpm at an angle of 80° for 1 minute and 50 seconds using BioComp gradient maker. Polysome sample lysate was added from the top of the tube and run on ultracentrifuge using an SW41 rotor (Beckman) at 36,000 rpm for 2 hours at 4° C.

### Polysome lysate preparation and fractionation

After completion of the experiment, 100µl of 10mg/mL cycloheximide was added to the media supernatant of 10 cm dish and kept at 37° C for 10 min. Samples were washed with 1X PBS (100µg/mL cycloheximide) followed by the addition of hypotonic buffer to the cell pellet (5 mM TRIS-HCl pH-7.5, 5 mM MgCl_2_ and 1.5 mM KCl). 3 minutes post incubation with hypotonic buffer, lysis buffer (5 mM TRIS-HCl pH-7.5, 5 mM MgCl_2_, 1.5 mM KCl, 100 µg/mL cycloheximide, 1mM DTT, 200 U/mL RNase in from Promega, 0.5% Sodium deoxycholate, 0.5% Triton X −100, 200µg t-RNA and 1X protease inhibitor cocktail) was added and kept in ice for 15 minutes. Later, KCl was added to adjust the final concentration to 150 mM and spun at 3000g for 8 min at 4° C. Supernatant was collected and run on ultracentrifugation as described before. After ultracentrifugation, samples were run on a polysome fractionator (BioComp) and absorbance for each drop was taken at 254 nm to generate the polysome profile of the samples. The collected samples were stored at −80° C and RNA was isolated later using TRI reagent for qRT-PCR.

### AlphaFold-based structural prediction

Full-length La protein (1-408aa) and cleaved La (1-295aa) were modelled using AlphaFold2-based [CollabFold](https://colab.research.google.com/github/sokrypton/ColabFold/blob/main/AlphaFold2.ipynb) platform(13). Default parameters were used for all predictions, and no structural templates were provided to avoid template bias. The Amber relaxation step was enabled to improve the stereochemical quality of the predicted structures. MSAs were generated automatically using MMseqs2 against UniRef and environmental sequence databases. Five model parameter sets were evaluated per model, and the highest-ranked model based on predicted Local Distance Difference Test (pLDDT) score was selected for downstream analysis.

### Hybrid structural assembly and refinement

To capture conformational heterogeneity of C-terminal IDR, structural ensembles were generated using IDP-GEN software (14). Selected IDR conformations were appended to cleaved La core (1-295aa) using the variable target function method (VTFM) followed by conjugate-gradient energy refinement and molecular dynamics with simulated annealing as implemented in MODELLER(15). Model selection was performed based on Discrete Optimized Protein Energy (DOPE) statistical potential score. The generated full-length models (IDR_full 1-10) were subsequently subjected to energy minimization prior to downstream analyses.

### Protein-Protein Docking and RNA-Protein Docking

Protein-Protein and RNA-Protein docking was performed using [HADDOCK](https://wenmr.science.uu.nl/haddock2.4/) (16, 17) to evaluate the effect of dimerization of La (full homo-dimer, cleaved-full hetero-dimer and cleaved homo-dimer) and interactions between La (IDR_full 1-10 and cleaved) and the SARS-CoV-2 5’UTR which was modelled using [RNAalifold](http://rna.tbi.univie.ac.at/cgi-bin/RNAWebSuite/RNAalifold.cgi). This was done using rigid-body energy minimization (it0), followed by semi-flexible simulated annealing refinement (it1) and final explicit-solvent refinement. Clustering was performed using the fraction of common contacts (FCC) metric with a 0.6 cutoff. Docked complexes were ranked according to the HADDOCK score, which integrates van der Waals, electrostatic, desolvation and restraint violation energies. The binding energy (ΔG) scores were obtained from Prodigy Webserver to analyse the binding affinities of the dimer complexes.

### Molecular Dynamics Simulations

All molecular dynamics (MD) simulations were performed using the GPU-enabled version of GROMACS. AMBER99SB-ILDN force field was employed to describe the protein systems, and TIP3P water model was used for solvation. The full-length La protein, cleaved La protein, and their respective RNA-bound complexes were individually placed in a cubic simulation box with a minimum distance of 1.0 nm from the box edges and solvated accordingly. Counter-ions were added to neutralize the systems. Energy minimization was carried out using the steepest descent algorithm until convergence. Equilibration was performed in two phases: NVT equilibration at 300 K using the Berendsen thermostat, followed by NPT equilibration at 1 bar pressure using the Parrinello - Rahman barostat. Position restraints were applied to heavy atoms during equilibration. Production MD simulations were conducted for 300 ns under periodic boundary conditions with a 2fs integration time step. Long-range electrostatic interactions were treated using the Particle Mesh Ewald (PME) method, and all covalent bonds involving hydrogen atoms were constrained using the LINCS algorithm. Trajectories were recorded every 10 ps for subsequent analysis. Structural stability and dynamic properties of full-length La, cleaved La, and their RNA-bound complexes were evaluated using RMSD, RMSF, radius of gyration (Rg), solvent-accessible surface area (SASA), and intermolecular interaction energy calculations as implemented in GROMACS. For MM-PBSA/MM-GBSA calculations, snapshots were extracted at regular intervals from the final 100 ns (200–300 ns) of the equilibrated trajectories to represent the converged binding ensemble. A total of evenly spaced frames was used for statistical averaging.

### Mass spectrometric analysis

Peptides were analyzed in LC-MS Orbitrap fusion (ThermoScientific). They were separated on a C-18 in-house column (15 cm length, 1.5µm particle size, internal diameter 150µm). Peptides were eluted with a gradient of 2%-90% acetonitrile in 0.1% formic acid and analyzed in positive-ion mode of the nano-electrospray ionization source. The mass spectrometer was operated in a data-dependent mode, automatically switching between MS and MS/MS acquisition. Automated MS data of peptides were acquired through Orbitrap mass analyzer between 350 m/z and 2000 m/z above the 5,000count threshold. The 20 most intense ions were selected for MS/MS acquisition using an ion trap mass analyzer. The ion charge states were +2 to +8. The higher-energy C-trap dissociation (HCD) fragmentation energy was adjusted to 30%. Raw data was analyzed using Proteome Discoverer V2.5 (ThermoScientific).

### Ethics and animal husbandry

The work plans for the animal experiment was assessed and approved by the Institute Biosafety Committee (IBSC) and the Institute Animal Ethical Committee (IAEC), and the experiment was conducted in compliance with CPCSEA (The Committee for the Purpose of Control and Supervision of Experiments on Animals) guidelines. Six- to eight-week-old female BALB/c mice were used in this study. The experimental mice were housed in individually ventilated cages (IVCs) with a 12-hour day/night light cycle, pellet diet, and unlimited access to water. In addition, the viral BSL-3 lab was maintained at 23±1^◦^C temperature and 50±5% relative humidity.

### Virus Infection experiments

In the virus BSL-3 laboratory, the experimental mice were randomly assigned to vehicle control and infected groups (n=5) after acclimating to IVC cages for seven days. A combination of ketamine (90 mg/kg/b.wt) and xylazine (4.5 mg/kg/b.wt) was administered intraperitoneally to sedate and anesthetize the mice. They were then intranasally infected with 10^5^ PFU of SARS-CoV-2 (MA-10 isolate) in 40 µL PBS. On the 5^th^ day of post infection, the mice were euthanized by anesthetizing the mice with isoflurane and followed by cervical dislocation. The lungs were then harvested for further experimental procedures.

## Results

### La protein binds to the GCAC sequence near initiator AUG of SARS-CoV-2 5’UTR

The length of SARS-CoV-2 5’ UTR is around 265 nucleotides (nt), which consists of five stem loop structures, conserved across variant of concerns (VoCs) (18). To investigate the binding profile of host RBPs with the 5’UTR of SARS-CoV-2, S10 lysates from HEK293T-ACE-2 were incubated with α-^32^P labelled 5’UTR (*in vitro* transcribed) followed by UV crosslinking and gel SDS-PAGE analysis. Phosphor imaging analysis showed binding of several RBPs of different sizes with the 5’UTR. Previously, we have reported two important RBPs-Human antigen R (HuR,35 kDa) and Polypyrimidine tract binding protein (PTB, 57 kDa), that interact with the 5’UTR of SARS-CoV-2 (19). Here, we observed another uncharacterized RBP around 52kDa (marked with an asterisk) which showed concentration dependent increase in binding (Fig 1A). To further characterize this RBP, we excised the band around 50 kDa and performed mass spectroscopy. Top 15 proteins (as per percentage of peptide coverage) were further analysed and identified one RBP, human La protein (in 50-55 kDa size range), along with few other host proteins (Fig 1B). The interaction was further characterized by direct and competitive UV crosslinking assays, using recombinant RBP La. HuR, a previously characterized RBP was used as a positive control for the experiment (Fig 1C) (19). Competitive UV crosslinking showed that 100-fold excess of self-RNA (unlabeled) could compete with α-^32^P labelled 5’UTR for binding with La protein. In contrast, Non-specific RNA (Nsp) did not show any competition (Fig 1D). Surface Plasmon Resonance (SPR) revealed high affinity binding of La protein with the 5’UTR (K_D_ 3.22 nM, K_a_= 1.07×10^6^M^-1^sec^-1^ and K_d_= 3.45×10^-3^sec^-1^) (Fig 1E).

**Fig 1.**
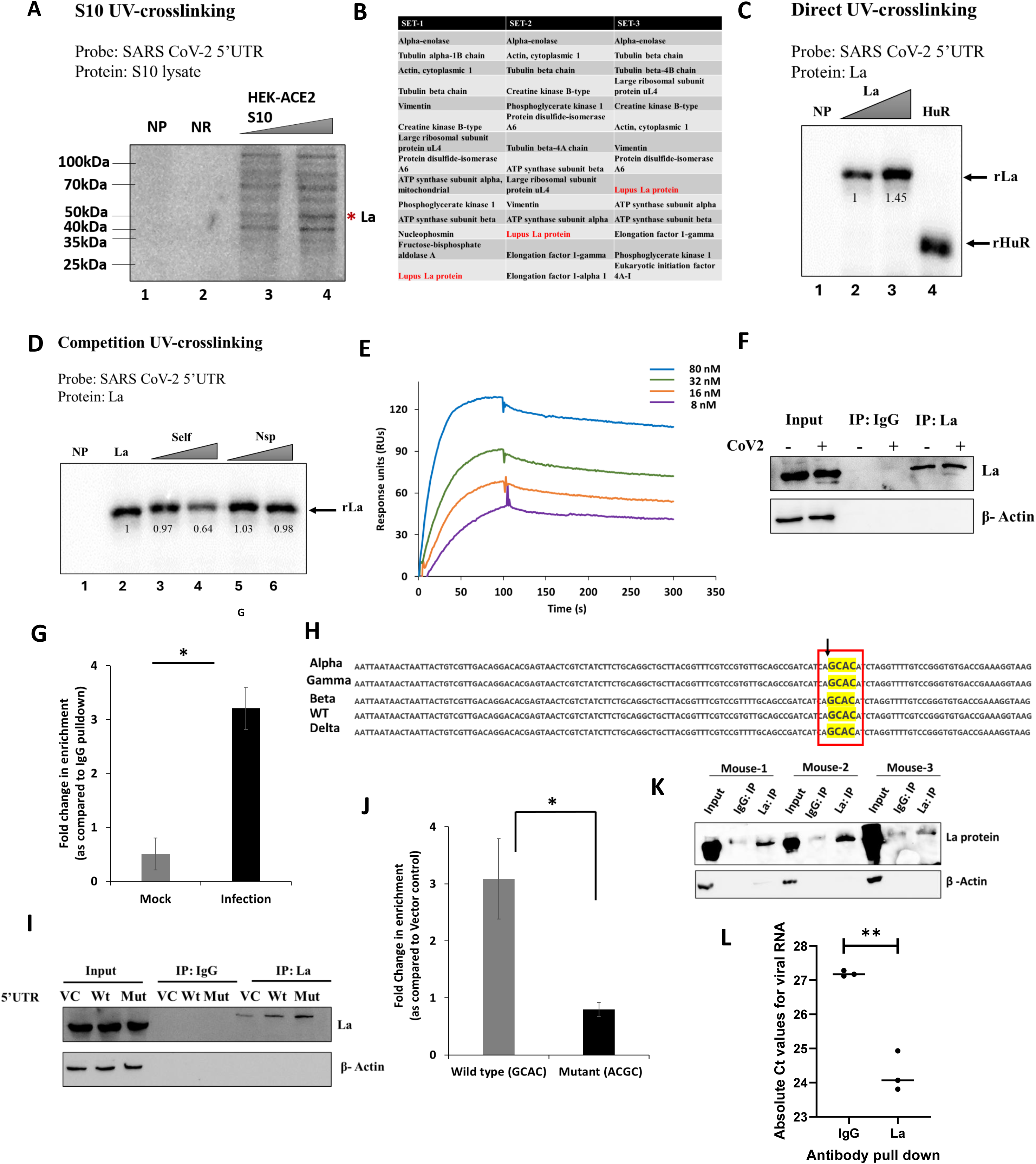
Lupus La protein binds with 5’UTR of SARS-CoV-2. (A) *in vitro* UV crosslinking of radio-labelled 5’UTR of SARS-CoV-2 with increasing concentration of S10 lysate of HEK-293T-ACE2 cells and Huh7 cells. The asterisk indicated unidentified RBP of interest (just beneath 50kDa). (B) Top 15 host proteins (as per % of peptide coverage) identified by mass-spectrometric analysis from the cut band beneath 50kDa size (3 sets). Lupus La protein was highlighted (red color) as only detected RBP. (C) *in vitro* UV crosslinking of recombinant La protein and radio-labelled 5’UTR of SARS-CoV-2. HuR was used as a positive control. (D) *in vitro* Competition UV crosslinking of recombinant La along with radio-labelled 5’UTR and unlabelled 5’UTR RNA. Unlabelled Nsp RNA was used as negative control. (E) Sensorgrams for indicated biotinylated SARS-CoV-2 5’UTR binding to recombinant La protein using SPR. Color denotes increasing concentration of La protein. (F) *ex vivo* interaction of La protein with SARS-CoV-2 infected HEK-293-ACE2 cells checked by immune-precipitation of La protein. Representative western blot suggested pulldown of La protein by anti-La antibody. IgG was used as negative control for the pull-down assay. (G) Enrichment of viral RNA with La protein upon infection was compared with mock. Association of viral RNA with La protein was normalized to IgG pulldown for each condition (N=3). (H) Multiple sequence alignment of 5’UTR of SARS-CoV-2 VoCs (WT, alpha, beta, gamma and delta). Conserved GCAC site was highlighted in yellow. (I) *ex vivo* interaction of La protein with SARS-CoV-2 Wild-type (Wt) and mutant (Mut) 5’UTR upon transfection in HEK-293-ACE2 cells. Representative western blot suggested pulldown of La protein by anti-La antibody. (J) Enrichment of viral Wt 5’UTR with La protein was compared with viral Mut 5’UTR. Association of 5’UTR RNA with La protein was normalized to IgG pulldown for each condition (N=3). (K) *in vivo* interaction of La protein with SARS-CoV-2 (MA-10 mice adapted strain) infected BALB/c mice checked by immunoprecipitation of La protein. Representative western blot suggested pulldown of La protein by anti-La antibody. IgG was used as negative control for the pull-down assay (n=3). (L) Association of viral RNA with La protein for individual mouse was plotted using absolute CT values and compared with absolute CT value of viral RNA association with IgG (negative control for pulldown assay) (n=3). Student’s t-test was used for statistical analysis. *=p<0.05, **=p<0.01, ***=p<0.001.

To confirm the binding of La protein with 5’UTR upon SARS-CoV-2 infection ex vivo, HEK-293T-ACE2 cells were infected with the wild-type strain of SARS-CoV-2 at MOI=1. Infected cells were harvested at 24 h.p.i and immunoprecipitated (using anti-La antibody) (Fig 1F). Quantitative real time PCR (RT-qPCR) from La pulled down fraction showed significant enrichment of virus RNA compared to IgG pulldown (Fig 1H). La protein was previously reported to interact with GCAC motif near iAUG of HCV 5’UTR RNA(12). Intriguingly, *in silico* analysis suggested coronaviruses across different species showed GCAC sequences in the 5’UTR and the 5’UTR of SARS-CoV-2 also contained two sets of GCAC sequences (Fig S1A & S1B). Multi-alignment of the 5’UTR showed that GCAC (227-230 nt) sequence motif proximal to the initiator AUG was conserved across alpha, gamma, beta, wild-type (WT) and Delta variant (Fig 1G). Site-directed mutagenesis (SDM) of 5’UTR, where GCAC (227-230 nt) was mutated to ACGC was used for further validation. The predicted secondary structure of wild type and the mutated 5’UTR, did not show significant change in the secondary structure (Fig S1C, S1D). To test the impact of mutation on 5’UTR binding inside the cells, wild-type and mutant 5’UTR RNA was transfected in HEK-ACE2 cells and 48h post transfection, immunoprecipitation was performed using anti-La antibody, followed by probing the enrichment of 5’UTR RNA. (Fig 1I). Wild-type 5’UTR showed significant enrichment along with La pulled down fraction compared to Mutant 5’UTR, suggesting that GCAC sequence near the initiator AUG is important for binding with La protein (Fig 1J). To further confirm the observation *in vivo*, in mouse infection model, BALB/c mice were infected with MA10 mice adapted strain of SARS-CoV-2 (Fig S1F). Lung tissue lysates were further processed to pull down mouse La protein (mouse La has almost 76% amino acid sequence similarity with human La) (Fig 1K, S1E). Results showed relatively more association of virus RNA with La protein (Absolute CT around 24) compared to IgG negative control (Absolute CT around 27) (Fig 1L). Results firmly establish that La protein specifically interacts with SARS-CoV-2 5’UTR RNA in the *in vitro, ex vivo* and *in vivo* experimental conditions.

### La protein differentially regulates viral RNA translation and replication

La protein binds near initiator AUG of 5’UTR of SARS-CoV-2, suggesting a possible role of La in initiation of viral RNA translation. To investigate its role in virus translation, we overexpressed La protein in HEK293T-ACE-2 cell line and 24 hours post transfection, the cells were infected with the SARS-CoV-2 virus at MOI=1 (Fig 2A). Cells were harvested 24h post infection, and polysome fractionation was performed (Fig 2B, 2C). Polysome profile of cells infected with virus La overexpression did not affect the overall distribution of monosome, and polysome peaks compared to mock infection (vector control vs La over expression). However, RT-qPCR data showed that association of viral RNA with the polysome was significantly increased upon overexpression of La protein (Fig 2D). Results suggest La protein might help recruiting virus RNA to polysome for active translation. Also, overexpression of La increased viral protein synthesis (Fig 2A), which reconfirmed our earlier observation.

**Fig 2.**
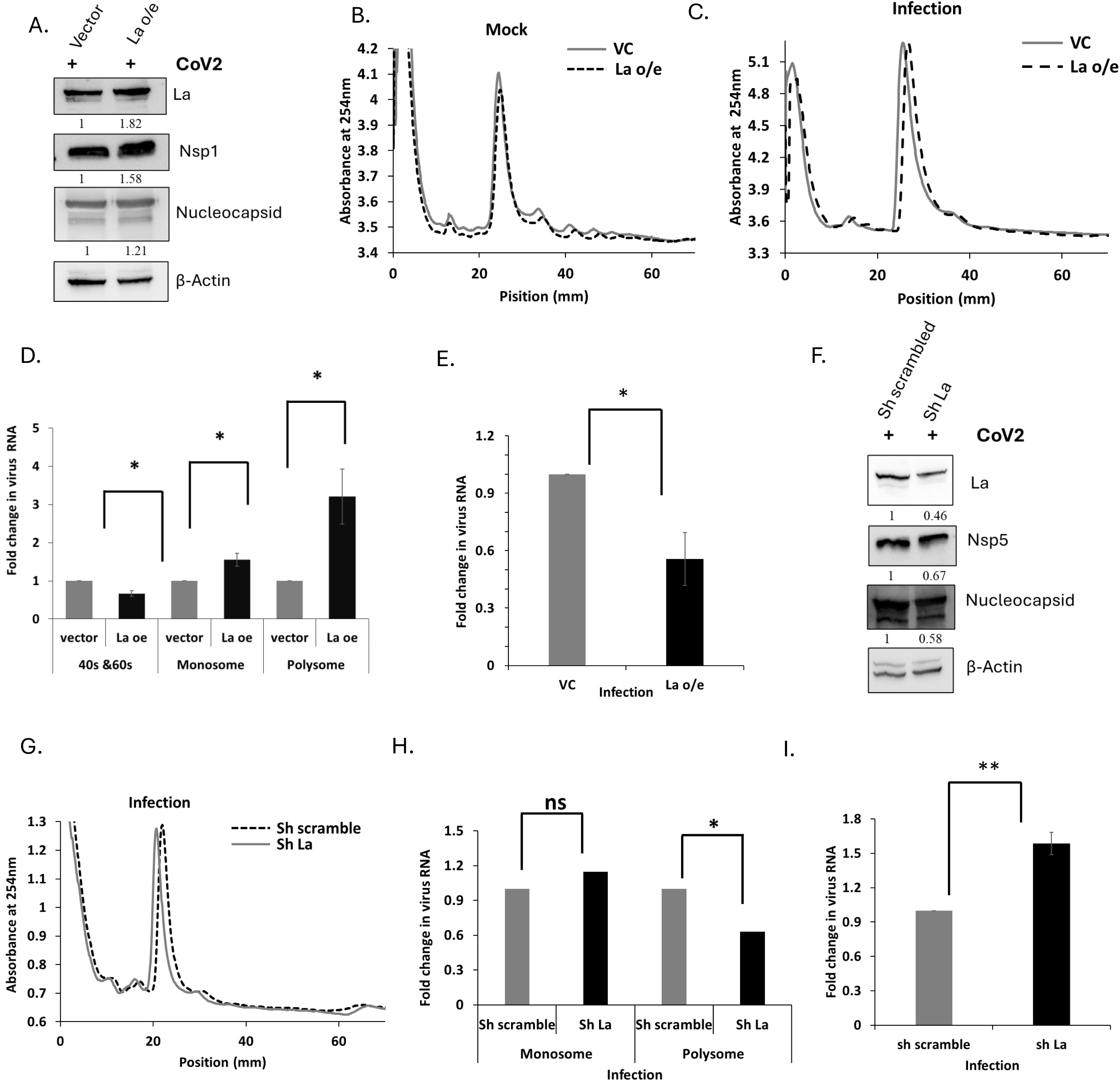
Host La protein differentially regulates virus translation and replication. (A) HEK-293T-ACE2 cells were transfected with pcDNA3.1-La overexpression construct and pcDNA3.1 (Vector control, VC) and followed by infected with SARS-CoV-2 at MOI=1. Representative western blot depicted overexpression of La protein and its impact on virus proteins (structural and non-structural proteins). (B, C) Representative graph depicted comparative polysome profile (40S, 60S, monosome and polysome) of VC and La overexpressed condition in uninfected (mock) cells and infected cells for polysome profile experiment. (D) RNA isolated from 40S, 60S, monosome and polysome fractions of infected cells and viral RNA was quantified by qRT-PCR. Expression of viral RNA from different fractions of La overexpression condition was compared to different fractions of vector control (VC) respectively (N=3). β-actin was used as housekeeping gene for all qRT-PCR data analysis. (E) RNA isolated from input fractions of polysome profiling experiment and qRT-PCR data was plotted. (F) HEK-293T-ACE2 cells were transfected with Sh La construct and Sh scramble and followed by infection with SARS-CoV-2 at MOI=1. Representative western blot depicted partial silencing of La protein and its impact on virus proteins (structural and non-structural proteins). (G) Representative graph depicted comparative polysome profile (40S, 60S, monosome and polysome) of infected cells post transfection with sh scramble and sh La. (H) RNA isolated from 40S, 60S, monosome and polysome fractions of infected cells and viral RNA were quantified by qRT-PCR. Expression of viral RNA from different fractions of partially silenced La condition was compared to different fractions of sh scramble respectively (N=3). (I) RNA isolated from input fractions of polysome profiling experiment and qRT-PCR data was plotted. Student’s t-test was used for statistical analysis. *=p<0.05, **=p<0.01, ***=p<0.001.

Structural proteins are synthesized by the sub genomic RNA while nonstructural proteins are produced by the genomic RNA of SARS-CoV-2. The 1-69 nt of the 5’UTR of genomic RNA serves as the 5’UTR of sub genomic RNA, that governs translation of structural proteins. Since La protein binds to GCAC motif at 227-230 nt, which is far away from the leader sequences (1-69nt), very little change in the production of structural proteins was observed, compared to non-structural proteins (Fig 2A). Intriguingly, virus RNA isolated from input fraction (infected lysates) of the polysome experiment showed significant reduction upon La overexpression, suggesting that La protein might negatively regulate virus replication (Fig 2E).

To further validate our findings, polysome fractionation was performed with lysates after partial silencing of La protein. For this purpose, HEK 293-ACE2 cells were transfected with La short hairpin RNA (shRNA) and 24 hours post transfection the cells were infected with the virus at MOI=1 (Fig 2F). Results suggest, partial silencing of La did not affect the overall distribution of monosomes and polysome peaks (Fig 2G). However, partial silencing of La protein resulted in a decreased association of viral RNA with the polysomes (Fig 2H). We further showed that partial silencing of La resulted in a decreased production of viral proteins (Fig 2F). However, Viral RNA isolated from the input fraction (infected lysates) used in the polysome experiment showed significant increase upon partial silencing of La protein, suggesting negative role of La protein in viral RNA replication. Results support our previous observations that La protein differentially regulates viral RNA translation and replication by enhancing translation while restricting RNA replication.

Since SARS-CoV-2 RNA mainly resides in cytoplasm of the cell upon infection, interaction between La protein and virus RNA could happen only in cytoplasm. In fact, subcellular fractionation (24 hours post infection) confirmed that cytoplasmic La levels remain unchanged upon infection (Fig S2A &S2B), that Results indicate that preexisting cytoplasmic La is sufficient to regulate virus translation and limit replication.

### La protein enhances cap independent translation of SARS-CoV-2 RNA

La protein could bind to the 5’UTR of SARS-CoV-2 and positively regulate virus translation. To investigate whether the effect was due to the binding of La protein with 5’UTR of SARS-CoV-2, we performed luciferase-based reporter assay. For this purpose, La overexpression construct was transfected in HEK293T-ACE2 cells and 24 hours post first transfection, another plasmid construct 5’UTR-CoV2 Renilla luciferase (Rluc) construct along with Firefly luciferase (Fluc, internal control) construct (Fig 3A) were transfected. No significant change in the relative Rluc values (normalized to Fluc) was observed upon overexpression of La protein compared to vector control (Fig 3B). Comparison of the SARS-CoV-2 (MOI=1) infected cell polysome profile with mock cells, showed a drastic induction in monosome peak and reduction in polysome peak upon infection (Fig 3C), suggesting virus mediated shut down of global translation. Interestingly, SARS-CoV-2 infection induced global translation shutdown, as evidenced by polysome collapse, was accompanied by a shift towards cap independent translation like different other lytic viruses (Fig S2C). Since virus relied mostly on cap independent translation upon repression of host cell cap dependent translation (4), we hypothesized that La could regulate cap independent translation of SARS-CoV-2.

**Fig 3.**
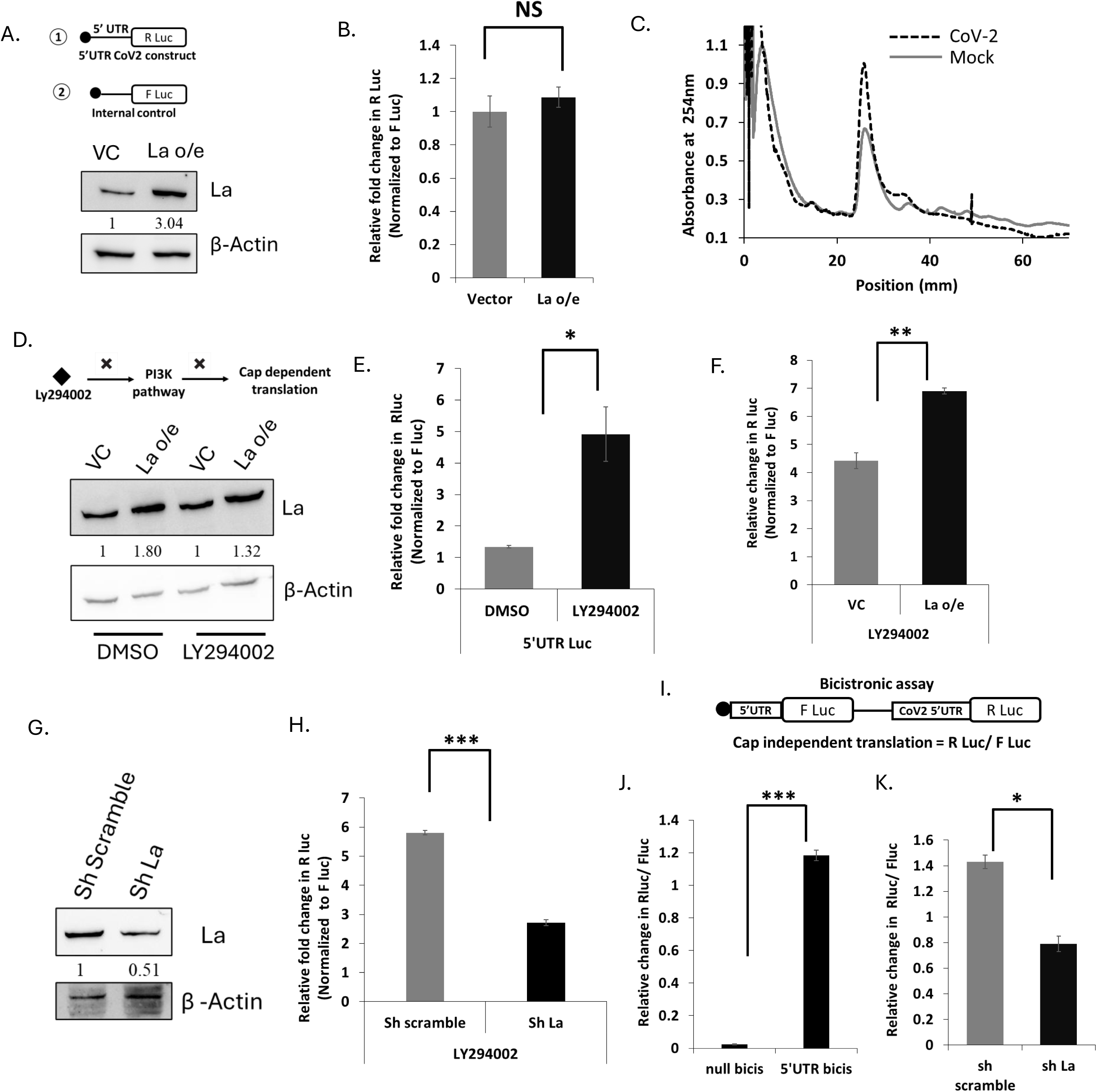
La protein regulated virus translation by cap independent mechanism. (A) Schematic of pcDNA3.1-5’UTR-Rluc construct and pcDNA3.1-Fluc construct. HEK-293T-ACE2 cells were transfected with pcDNA3.1-La overexpression construct and later transfected with luciferase construct. Representative western blot depicted overexpression of La protein. (B) Relative fold change in Rluc was plotted for vector control and La overexpression conditions. Absolute Rluc values were normalized to Fluc (transfection internal control). (C) Representative graph depicted comparative polysome profiles (40S, 60S, monosome and polysome) of mock and infected HEK-293T-ACE2 cells for polysome profile experiment. (D) Schematic represents mechanism of action of LY294002, PI3K inhibitor. HEK-293T-ACE2 cells were transfected with pcDNA3.1-La overexpression construct and later transfected with luciferase construct and LY294002 was treated post second transfection. DMSO was used as control. Representative western blot depicted overexpression of La protein upon DMSO and LY294002 treated condition. (E) Relative fold change in Rluc was plotted upon DMSO and LY294002 treated condition. Absolute Rluc values were normalized to Fluc. (F) Relative fold change in Rluc was plotted upon vector and La overexpression condition (LY294002 was treated in both condition). (G) HEK-293T-ACE2 cells were transfected with sh La construct and later transfected with luciferase constructs. Representative western blot depicted partial silencing of La protein. (H) Relative fold change in Rluc was plotted upon sh scramble and sh La overexpression condition (LY294002 was treated in both condition). (I) Schematic representation of SARS-CoV-2 bicistronic construct. Ratio of Rluc and Fluc depicts cap independent translation. (J) Relative fold change in Rluc/Fluc was measured for null bicis construct (control) and SARS-CoV-2 5’UTR bicis construct. (K) Relative fold change in Rluc/Fluc was measured for SARS-CoV-2 5’UTR bicis construct upon partial silencing of La protein. Student’s t-test was used for statistical analysis. *=p<0.05, **=p<0.01, ***=p<0.001 (N=3).

To further validate our hypothesis, similar experiment was repeated post treatment with LY294002 (an inhibitor of PI3K pathway, which inhibits cap dependent translation). Briefly, La overexpression construct was transfected and 18 hours post transfection, cells were treated with 25µM of LY294002. After 4 hours of drug treatment, 5’UTR-CoV2 Rluc and Fluc (internal control) constructs were transfected (Fig 3D). Results showed a significant increase in relative Rluc values upon LY294002 treatment compared to DMSO control, suggesting an increase in cap independent translation (Fig 3E). La overexpression showed further increase in cap independent translation compared to vector control under LY294002 treated conditions (Fig 3F). In parallel, we also performed partial silencing of La protein in cells using shLa construct followed by treatment of LY294002 (Fig 3G). 4 hours post drug treatment, we transfected 5’UTR-CoV2 Rluc and Fluc constructs. Partial silencing of La showed significant decrease in cap independent translation (Fig 3H). To further validate these observations, bicistronic constructs with luciferase reporter genes were used, where 5’UTR of SARS-CoV-2 is flanked in between Fluc and Rluc (Fig 3I). Ratio of Rluc and Fluc (Rluc/Fluc) indicates a measure of cap independent translation capacity of 5’UTR of SARS-CoV2 (Fig 3J). Partial silencing of La protein could significantly decrease the ratio of Rluc to Fluc compared to vector control (Fig 3K), suggesting a reduction in cap independent translation, which further strengthened our earlier observations with the drug (LY294002) treated cells. Results clearly demonstrated that La specifically enhances cap-independent translation mediated by the SARS-CoV-2 5’UTR RNA and strongly establishes La as a positive regulator of SARS-CoV-2 cap-independent translation. Previous reports on SARS-CoV2 and our experimental evidences further suggested that cap independent translation is more pronounced in Genomic 5’UTR rather than subgenomic 5’UTR of SARS-CoV-2 (Fig S2D). Since, La binds with Genomic 5’UTR of SARS-CoV-2, it further explained its regulation on cap independent translation of Genomic 5’UTR.

### Nsp5 protease of virus proteolytically cleaves La protein upon infection

SARS-CoV2 infection is known to change the expression profile of different host factors including RBPs (19). To check whether abundance of La protein is altered upon virus infection, La protein level upon SARS-CoV-2 infection (MOI=1) was probed by western blot analysis. Interestingly, upon infection, an additional lower molecular weight band along with expected La protein band (52 kDa) was observed (Fig 4A & 4B). To investigate whether this additional band is due to the non-specific detection by anti La antibody (EPR-6570, Abcam), we further verified the detection of this band by using another anti La antibody (NBP1-48802, Novus biologicals) (Fig S3A). Since different mRNA isoforms of SSB gene produce only full length La protein of 52kDa, we hypothesized that additional band could be partially cleaved La protein, generated by caspase cleavage or virus protease mediated cleavage. SARS-CoV2 virus encoded two proteases Nsp5 and Nsp3 (20). To investigate whether one of these virus proteases could cleave La protein, mammalian expression constructs encoding either Nsp5 and Nsp3 were transfected and 48h post transfection La protein was probed by western blot analysis. The additional band showed up only upon ectopic expression of Nsp5 and not by Nsp3 or mock condition, suggesting that Nsp5 protease could cleave La protein (Fig 4C). It is reported that Nsp5 protease could induce apoptosis through Bcl-II axis (21). To confirm that the cleavage of La protein is due to catalytic activity of Nsp5, we transfected Nsp5 wild-type and Nsp5 Catalytic dead mutant (C145A) and harvested cells at 48h post transfection. Results showed that Nsp5 wild-type could cleave La protein but not the Nsp5 catalytic dead mutant, suggesting that catalytic activity of Nsp5 protease is necessary for cleavage of La protein (Fig 4D). Further, overexpression with increasing concentration (100ng, 150ng, 200ng) of Nsp5 wild-type (WT) showed dose dependent increase in cleavage percentage (Fig 4E). These observations strengthened our earlier results.

**Fig 4.**
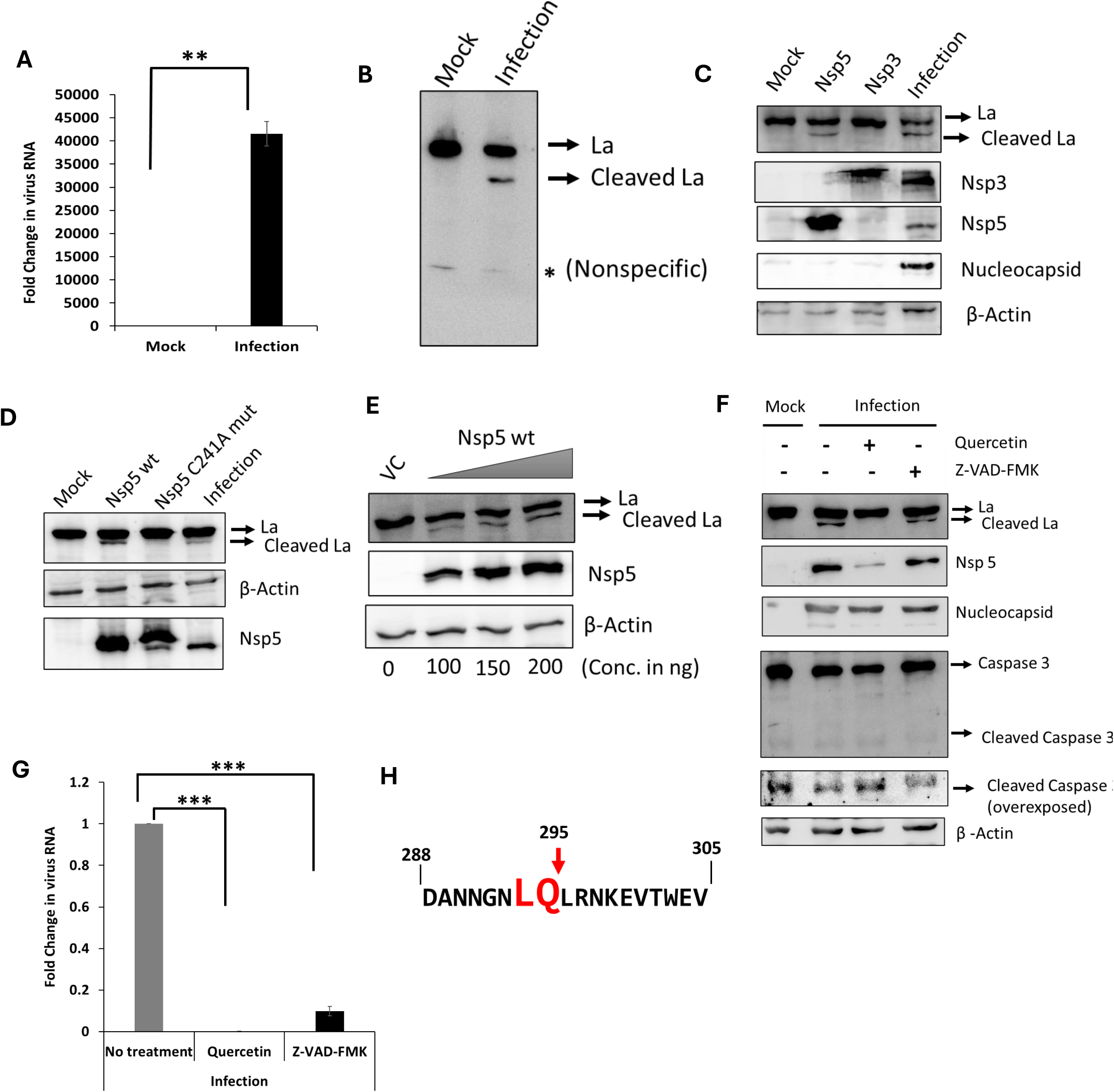
Nsp5 protease of SARS-CoV-2 proteolytically cleaves host La protein upon infection. (A) HEK-293T-ACE2 cells were infected with SARS-CoV-2 at MOI=1. Expression of Viral RNA was quantified by qRT-PCR. (B) Representative western blot depicted expression level of La protein 24h post virus infection. Additional band upon infection predicted as cleaved La.(C) HEK-293T-ACE2 cells were transfected with SARS-CoV-2 Nsp5 and Nsp3 overexpression construct. Representative western blot depicted effect of Nsp5 and Nsp3 overexpression on cleavage of La protein. SARS-CoV-2 infected cell lysate was used as positive control. (D) HEK-293T-ACE2 cells were transfected with Wild-type Nsp5 (Nsp5 Wt) and catalytically dead mutant Nsp5 (Nsp5 C145A Mut) overexpression construct. Representative western blot depicted effect of Nsp5 Wt and Nsp5 C145A Mut overexpression on cleavage of La protein. SARS-CoV-2 infected cell lysate was used as positive control. (E) HEK-293T-ACE2 cells were transfected with increasing concentration (100ng, 150ng and 250ng) of Nsp5 Wt overexpression construct. Representative western blot depicted effect of concentration dependent Nsp5 Wt overexpression on cleavage of La protein. (F) HEK-293T-ACE2 cells were infected with SARS-CoV-2 at MOI=1 and 1 h.p.i, cells were treated with quercetin (inhibitor of Nsp5 protease) and Z-VAD-FMK (pan caspase inhibitor). DMSO was used as control for drug treatment. Representative western blot depicted effect of quercetin and Z-VAD-FMK on cleavage of La protein. (G) Representative graph depicted effect of quercetin and Z-VAD-FMK on viral RNA. Viral RNA quantified by qRT-PCR. (H) Amino acid sequence of La protein depicted recognition site of Nsp5 protease. Student’s t-test was used for statistical analysis. *=p<0.05, **=p<0.01, ***=p<0.001 (N=3).

It is also reported previously that La protein is cleaved during apoptosis due to caspase activity (22, 23). To rule out the possibility of caspase mediated cleavage of La protein upon SARS-CoV-2 infection, HEK 293T-ACE2 infected cells were infected with SARS-CoV-2 virus at MOI=1 followed by treatment with quercetin (Nsp5 inhibitor) and Z-VAD-FMK (pan caspase inhibitor) after 1h postinfection. Results showed that Nsp5 inhibitor, quercetin could prevent cleavage of La protein by restricting catalytic activity of Nsp5 protease but the pan caspase inhibitor, Z-VAD-FMK, could not restrict the cleavage (Fig 4F). Also, induction of apoptosis (activated cleaved caspase 3) was not observed until 24 hours post infection, suggesting cleavage of La is likely due to catalytic activity of Nsp5 protease only (Fig 4F).

Additionally, to check whether other variants of concern (VoCs) of SARS-CoV-2 could cleave La protein, HEK-293T-ACE2 cells were infected with Delta and Omicron VoCs (at MOI=1) and observed similar cleavage of La protein 24 hours post infection (Fig S3B). Also, quercetin and Z-VAD-FMK could inhibit SARS-CoV-2 RNA level (Fig 4G).

SARS-CoV-2 Nsp5 protease is known to recognize its substrate by presence of the LQ motif (24). Incidentally, the amino acid sequence analysis of La protein revealed the presence of a conserved LQ motif at 295 aa positions, which further confirmed the mechanism of action by which Nsp5 cleaved La protein (Fig 4H) suggesting host La protein as a direct substrate of the viral Nsp5 protease.

### Cleavage of La protein impairs its binding with viral 5’UTR and reduced translation

To investigate the functional consequences of La cleavage, Nsp5 WT and Nsp5 dead mutant were over expressed separately along with the 5’UTR-CoV-2-Rluc construct. 48 hours post transfection, association of La protein with the 5’UTR RNA was probed by immunoprecipitation of La protein (using anti La antibody) (Fig 5A). Our RT-qPCR data showed reduced association of viral 5’UTR with La protein upon overexpression of Nsp5-WT compared to Nsp5 dead mutant. (Fig 5B), suggesting cleaved La (due to Nsp5 WT overexpression) showed reduced interaction with 5’UTR of SARS-CoV-2.

**Fig 5.**
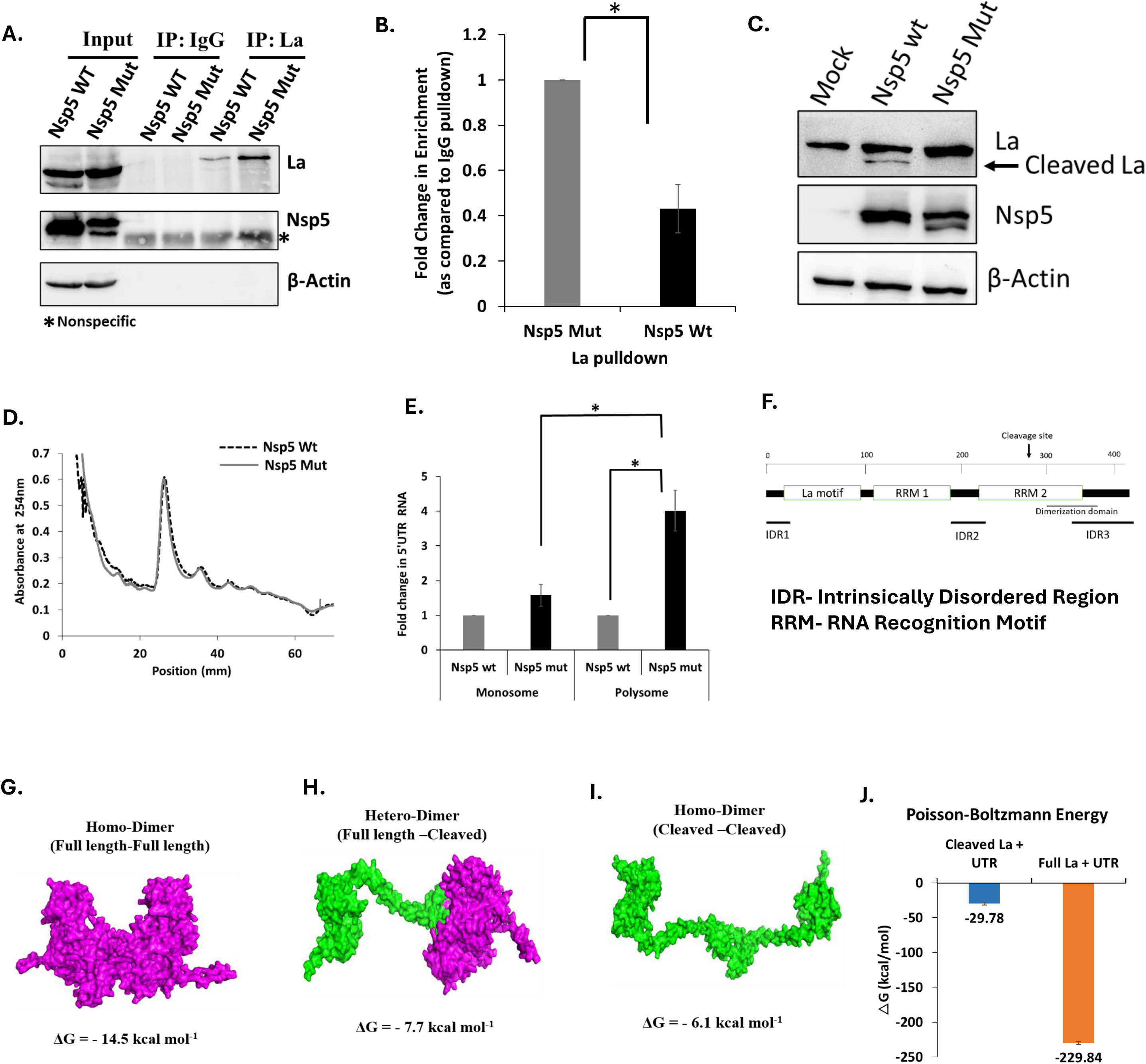
Cleaved La showed reduction in virus translation due to reduced interaction with 5’UTR of SARS-CoV-2. (A) HEK-293T-ACE2 cells were transfected with Nsp5 Wt and Nsp5 C145A Mut overexpression construct. 5’UTR-CoV-2-Rluc was transfected for both conditions. 48h post transfection, La protein was immunoprecipitated using anti-La antibody. Representative western blot depicted pulldown of La protein Upon Nsp Wt and Nsp5 C145A Mut overexpression condition. (B) Enrichment of 5’UTR with La protein upon Nsp5 C145A Mut overexpression was compared with Nsp5 Wt overexpression by qRT-PCR. Association of 5’UTR with La protein was normalized to IgG pulldown for each condition (N=3). (C) HEK-293T-ACE2 cells were transfected with Nsp5 Wt and Nsp5 C145A Mut overexpression construct. 5’UTR-CoV-2-Rluc was transfected for both conditions. 48h post transfection, samples were harvested for polysome profiling. Representative western blot depicted overexpression of Nsp Wt and Nsp5 C145A Mut and its effect on cleavage of La protein. (D) Representative graph depicted comparative polysome profile (40s, 60s, monosome and polysome) of Nsp5 Wt and Nsp5 C145A Mut overexpression condition. (E) RNA isolated from 40S, 60S, monosome and polysome fractions for both conditions (Nsp Wt and Nsp5 C145A Mut overexpression) were quantified by qRT-PCR. (F) Schematic diagram of La protein. (G) Represented diagram depicted dimerization domain guided specific docking between the stitched and simulated structures of two full length La protein monomer. (H) Represented diagram depicted blind docking between simulated structure of full-length La and cleaved La monomer. (I) Represented diagram depicted blind docking between simulated structures of two cleaved La monomer. (J) MM/GBSA and MM/PBSA binding free energy calculations demonstrate markedly stronger binding of full-length La to the SARS-CoV-2 5′UTR (ΔG = −229.84 ± 2.36 kcal/mol) compared to the cleaved protein (ΔG = −29.78 ± 1.85 kcal/mol). Student’s t-test was used for statistical analysis. *=p<0.05, **=p<0.01, ***=p<0.001 (N=3).

To understand the impact of decreased association of 5’UTR with La protein, polysome profiling experiment was performed by transfecting Nsp5 WT and Nsp5 dead mutant along with 5’ UTR-CoV2-Rluc construct in HEK293T-ACE2 (Fig 5C). No significant change was observed in the overall polysome profile of cells overexpressed with Nsp5 WT or Nsp5 dead mutant (Fig 5D). However, RNA isolated from monosome and polysome fractions showed increased association of virus RNA in the polysome cells overexpressing Nsp5 dead mutant compared to Nsp5 WT. (Fig 5E). Results confirmed our hypothesis that cleavage of La resulted in reduced binding with the viral 5’UTR, concomitant decrease in polysome recruitment of viral RNA, resulting in reduced viral RNA translation.

Earlier, it has been reported that dimerization of La protein is essential for promoting translation of Polio and Human immunodeficiency virus (HIV) (30). Since dimerization domain of La protein resides between 293-348 amino acids (Fig. 5F), and these residues are absent in the cleaved La variant (residues 1–295), which suggests that the truncated form may be defective in dimer formation. Since La dimerization is required for efficient IRES-mediated translation, we investigated whether Nsp5-mediated cleavage compromises this property. Guided docking showed that full-length La formed the most stable homodimer (ΔG = −14.5 kcal/mol) whereas full length/cleaved heterodimers (ΔG = −7.7 kcal/mol) and cleaved homodimers (ΔG = −6.1 kcal/mol) were markedly less stable, consistent with the loss of the dimerization domain (Fig. 5G-I; Fig 5SD). Along with dimerization domain, the loss of C-terminal intrinsically disordered region (IDR) upon cleavage suggested a potential role in RNA recognition, promoting further structural investigation (Fig. 5F). PONDR identified an IDR spanning residues 369-408, and stitching of representative IDR conformers onto the cleaved core generated full-length La models with substantially improved RNA-binding properties. Among these, IDR model 4 showed the strongest interaction with the SARS-CoV-2 5′UTR, exhibiting a markedly improved HADDOCK score, enhanced electrostatic interactions and a larger binding interface than cleaved La (Supplementary Fig. 5A-C). Molecular dynamics further demonstrated that the full-length La-RNA complex remained more stable than the cleaved complex, with lower backbone deviation, a more favorable free-energy landscape and substantially stronger MM/GBSA and MM/PBSA binding energies (Fig. 5SE-SJ; Fig. 5J). Together, these findings indicate that Nsp5-mediated cleavage simultaneously disrupts La dimerization and removes the positively charged, flexible IDR that stabilizes RRM2-mediated RNA recognition, thereby weakening La-5′UTR interactions required to support viral translation.

### Cleavage of La protein drives switching translation to replication

Till now we have shown that La protein positively regulated cap independent translation of viral RNA and negatively regulated viral RNA replication upon binding with 5’UTR of SARS-CoV-2. Also, one of the non-structural proteins, Nsp5 cleaved La protein. Intriguingly, cleaved La showed reduction in interaction with 5’UTR thus resulted in decreased translation. To further understand the time course of these events upon virus infection, we performed time kinetics after infecting HEK-293T-ACE2 cells with SARS-CoV-2 virus (MOI=1). It was observed that the non-structural proteins first showed up at 6 hours post infection and gradually increased synthesis 12 hours post infection (h.p.i). Whereas cleavage of La protein was observed at 18 h.p.i and gradually increased to 24 h.p.i (Fig 6A). The viral RNA estimation at different time-points revealed that initially at 6 hours and 12 h.p.i, virus RNA level did not show much replication but from 12 hours to 18 hours, there was an increase in replication (Fig 6B). These observations suggest that initially (6 to 12 h.p.i) SARS-CoV2 RNA translated to produce non-structural proteins, where La is one of the important RBPs that assisted virus translation in a cap independent manner. At later time point when La protein was partially cleaved by one of the non-structural proteins, Nsp5, it gradually switched from translation to replication.

**Fig 6.**
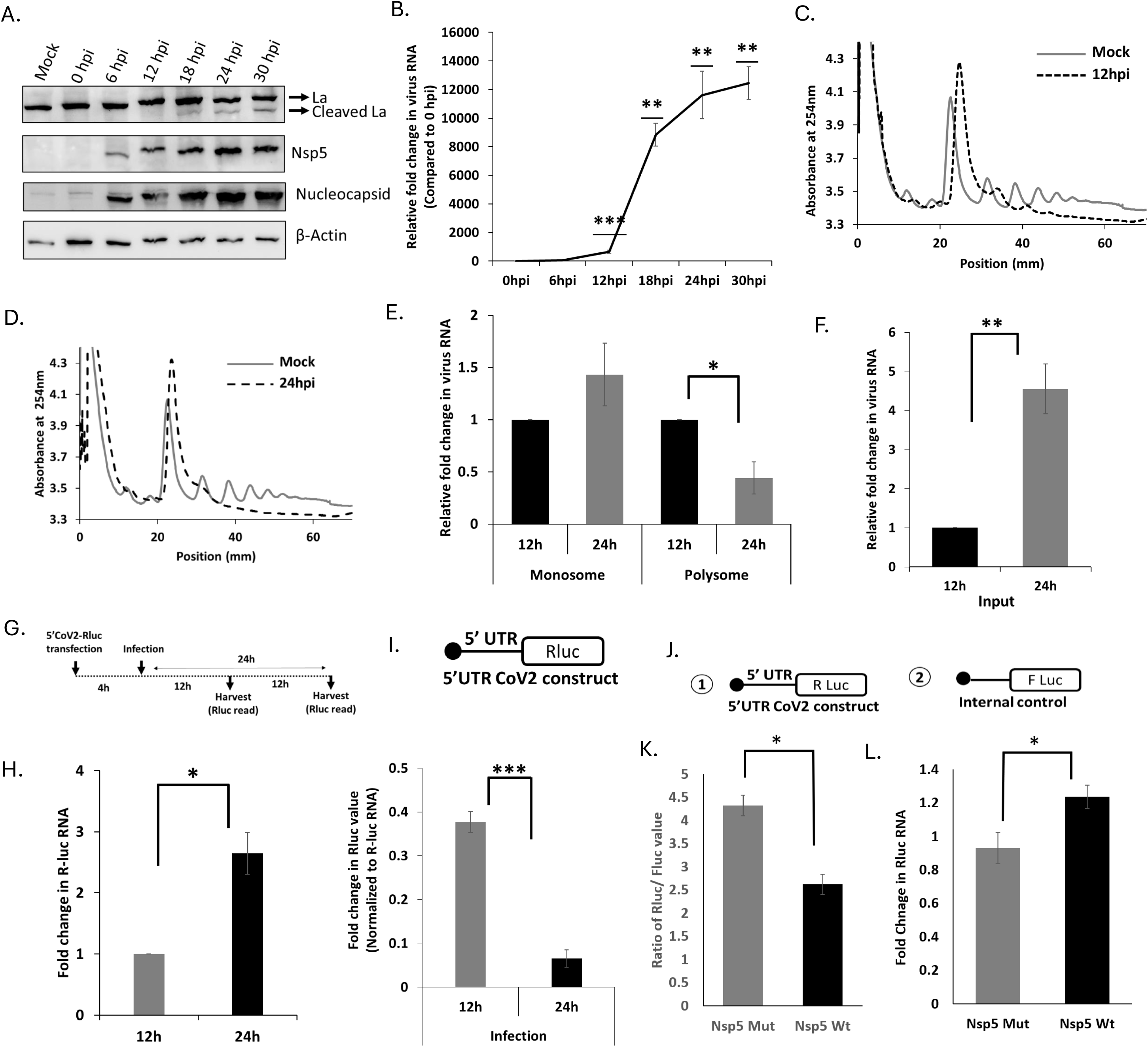
Cleavage of La protein assisted in switching translation to replication in SARS-CoV2. (A) HEK-293T-ACE2 cells were infected with SARS-CoV-2 at MOI=1 and samples were harvested at different timepoints (0h, 6h, 12h, 18h, 24h and 30h). Representative western blot depicted time kinetics of La protein expression and cleavage. (B) Representative graph depicted viral RNA abundance in different timepoints (0h, 6h, 12h, 18h, 24h and 30h) post infection. (C) Representative graph depicted comparative polysome profile (40s, 60s, monosome and polysome) of mock (uninfected) vs Infection (12h). (D) Representative graph depicted comparative polysome profile (40S, 60S, monosome and polysome) of mock (uninfected) vs Infection (24h). (E) RNA isolated from monosome and polysome fractions of infected samples harvested at 12h and 24h and viral RNA was quantified by qRT-PCR. (F) RNA isolated from input fraction (total RNA) of polysome profiled experiment and qRT-PCR data were plotted. (G) Schematic represents experimental timepoints. HEK-293T-ACE2 cells were transfected with 5’UTR-CoV-2 Rluc and 4h post transfection, cells were infected at MOI=1. (H) Representative graph depicted fold change in Rluc RNA at 12h and 24h post SARS-CoV-2 infection. (I) Representative graph depicted fold change in Rluc value at 12h and 24h post SARS-CoV-2 infection. Rluc value was normalized with Rluc RNA level. (J) Schematic of pcDNA3.1-5’UTR-Rluc construct and pcDNA3.1-Fluc construct. (K) Relative fold change in Rluc was plotted for Nsp5 Wt and Nsp5 Mut overexpression conditions. Absolute Rluc values were normalized to Fluc (transfection internal control). (L) Representative graph depicted fold change in Rluc RNA upon Nsp5 Wt overexpression compared to Nsp5 Mut. Student’s t-test was used for statistical analysis. *=p<0.05, **=p<0.01, ***=p<0.001 (N=3).

To further investigate this, we performed polysome profiling at 12 hours (when La protein was intact) and 24 hours (La protein was partially cleaved). We observed that at 12 hours, there was decreased polysome peak (reduction in disome and trisome peak) in virus infected cells compared to mock cells, suggesting virus infection led to partial shutdown in global translation (Fig 6C). However, at 24 h.p.i, virus infection led to complete shutdown in global translation (disome and trisome peaks were not visible) (Fig 6D). We showed that from 12 hours to 24 hours, there was a reduction in viral RNA in the polysome fraction, suggesting that virus translation was reduced from 12 to 24 h.p.i (Fig 6E). However, input RNA (total RNA), showed increase in virus RNA between 12 to 24 h.p.i, suggesting increase in replication (Fig 6F). Thus, it appears that virus RNA switched translation to replication at 24 h.p.i, which aligns with the timeline of cleavage of La protein by Nsp5.

To further confirm our hypothesis, luciferase-based reporter assays were performed in the presence of virus infection. 5’UTR-Rluc construct was transfected in HEK-293T-ACE2 and 4h post transfection, cells were infected with SARS-Co2 at MOI=1 (Fig 6G). Results showed that Rluc RNA levels increased at 24 h.p. i compared to 12 h.p.i though Rluc values (Normalized to Rluc RNA) reduced at 24 h.p.i compared to 12 h.p.i(Fig 6H, 6I). To further establish Nsp5 protease mediated switch, Nsp5 Wt and Mut constructs were overexpressed and 14h post first transfection, 5’UTR-Rluc construct was transfected (Fig 6J). Nsp5 Mut condition showed higher Rluc value (normalized to Fluc control) compared to Nsp5 Wt condition (Fig 6K), however subtle but significant increase in Rluc RNA was observed in Nsp Wt condition compared to Nsp5 Mut condition (Fig 6L). These observations indicate that La supports early translation whereas the cleavage mediated by viral Nsp5 promotes shift towards replication by freeing viral RNA from translation activity.

## Discussion

Host factors play essential role in regulating the lifecycle of different viruses at multiple stages of infection. In particular, host RBPs modulate virus replication and translation by interacting with viral RNA and proteins (26–28). Although several RBPs have been identified through high-throughput studies following SARS-CoV-2 infection, only a limited number of RBPs are functionally characterized. In this study, we showed La protein interacts specifically with the GCAC sequence near the iAUG of the 5’UTR of SARS-CoV2 upon infection. Previously, it was reported that La protein could interact with GCAC sequence at HCV 5’UTR, suggesting La protein binding site is conserved in few other viruses. Multi-alignment showed that not only GCAC site was conserved across different VoCs of SARS-CoV-2, but GCAC sites were present across different coronaviruses. Evolutionary conservation of GCAC sites suggested its importance for coronaviruses. Functional assays revealed that La protein negatively regulates virus replication and positively regulates the cap independent translation in virus. Interestingly, we observed that La preferentially promotes the early production of non-structural proteins translated from full length genomic RNA of SARS-CoV-2. Earlier, it was reported that full length 5’UTR of SARS-CoV-2 facilitates translation through cap independent mechanism but not sub genomic 5’UTR upon global translation shut down. The preferential effect of the La protein on non-structural protein can be explained by its regulation of viral cap-independent translation through binding to the full-length 5′ UTR. While La protein has previously been implicated in regulating IRES-mediated translation in Hepatitis C virus, Poliovirus, and Coxsackievirus B3, our study uncovers a novel function of La protein in controlling the differential expression of structural and non-structural proteins via cap-independent translation.

Though La protein drives cap independent translation of SARS-CoV-2 upon early infection, this protein is partially cleaved by Nsp5 protease of the virus. Nsp5 protease has been shown to cleave several host proteins, including histone deacetylases, innate immune regulators, and tRNA methyltransferases. Here, we report for the first time that Nsp5 also cleaves the RNA-binding host protein La, identifying a novel target of SARS-CoV-2 protease activity. We have further shown that partial cleavage of La results in reduction in binding with 5’UTR of SARS-CoV-2 and ultimately results in reduction in virus translation. It was previously reported that dimerization of La protein enhances virus translation (30). Given that La dimerization enhances viral RNA translation by stabilizing ribonucleoprotein complexes and promoting efficient ribosome recruitment, cleavage imposes a dual inhibitory effect: weakening RNA affinity and diminishing formation of translation-competent dimers and thereby mechanistically explaining the reduced polysome association and attenuated viral translation observed in the presence of the truncated protein. Our time kinetics experiments indicate that between 18-24 h.p.i, La protein undergoes proteolytic cleavage by the viral protease Nsp5. This cleavage correlates with reduced Viral RNA translation and enhanced replication at 24 h.p.i. Similar pattern of translation and replication was observed via luciferase-based reporter assay. Since we observed earlier that La protein negatively regulates replication and positively regulates translation, these time kinetics clearly suggested cleavage of La protein could be the key driver for virus translation to replication switch. Our reporter-based assay upon Nsp5 wild type and catalytic dead mutant expression further proved that cleavage of La regulated such switch. Since replication of virus RNA occurs in double membrane vesicle and translation occurs in open cytoplasm, same template RNA might not be used for replication and translation at same time. Taken together, our working model suggested that when la protein gets cleaved, a large pool of positive strand RNA of SARS-CoV-2 were no longer engaged with the polysomes and may instead be redirected toward the formation of replication complex. Interestingly, different reports suggested that RBPs could regulate translation to replication switch of viruses via displacing another RBP from virus UTRs. Since multiple RBPs could bind with SARS-CoV-2 5’UTR, La protein might drive the switch from translation to replication by similar mechanism as well. In summary, our study reveals a dynamic interplay between the host RNA-binding protein La and the SARS-CoV-2 protease Nsp5, that governs the switch from viral translation to replication during virus infection. Consistent with our findings on La protein and SARS-CoV-2, previous studies have also implicated La protein, as a critical host factor in life cycles of several other viruses (9, 10, 12) underscoring its broader relevance in viral RNA regulation. Taken together, these observations suggest that La could be a potential target for developing therapeutics to act against SARS-CoV-2. Notably, peptides such as LAP and LAR2C previously shown to play important role as inhibitor of virus translation by acting against La (31, 32). *In vitro* and *in vivo* studies evaluating the efficacy of these peptides in the context of SARS-CoV2 infection may provide new avenues for therapeutic intervention in future.

## Data availability

The datasets used in the current study are available within the manuscript and the Supplementary files.

## Acknowledgements

SD acknowledges the J.C. Bose Grant from ANRF, Department of Biotechnology (DBT), India, for research support. This study was also supported by Department of Biotechnology (DBT), Government of India, Indo-Swiss project, DBT-IISc partnership program, DST Fund for Improvement of Science and Technology Infrastructure (DST-FIST) level II infrastructure, and the University Grants Commission Centre of Advanced Studies. RS is supported by the Prime Minister Research Fellowship (PMRF) and contingency. We recognize the SPR facility at the Biological Science division at IISc. We acknowledge BEI resources, NIAID, NIH for providing the SARS-CoV-2 virus strains. We acknowledge Dr. Anup Majumder, NIBMG, Kalyani for Nsp5 wild type and Nsp5 (C241U) mutant overexpression constructs. We acknowledge Dr, Shashank Tripathi for Nsp3 overexpression constructs. We acknowledge Dr. Milan Surjit, Translational Health Science and Technology Institute (THSTI) for pCBB 5’-UTR constructs. We acknowledge viral BSL3 facility at CIDR, IISc supported by DBT-BIRAC and Crypto-Relief foundation for maintaining SARS-CoV-2 virus stocks.

## Author contributions

Conceptualization, R.S and S.D.; Methodology, R.S, G.G, S.P, H.R, S.V, P.K.G, and R.S.R; Investigation, R.S and S.D.; Resources, S.D.; Writing-Original, R.S and S.D.; Writing-Review and editing, R.S., G.G, and S.D.; Visualization, R.S. and S.D.; Supervision, S.D.; Funding acquisition, S.D.

---

**Fig S1 Prediction of La binding site across 5’UTRs of coronaviruses (**A) Multi-alignment of 5’UTRs across different coronaviruses depicted conservation of GCAC sites. (B) Secondary structure of 5’UTR of SARS-Cov-2 marked (red box) with putative La binding sites. (C) mfold predicted secondary structure of 5’UTR of SARS-CoV-2 containing wild type GCAC site. (D) mfold predicted secondary structure of 5’UTR of SARS-CoV-2 containing mutant ACGC site. (E) Multi-alignment of mouse La and human La protein sequence. (F) Representative image depicted histopathology of lung tissue of PBS control and MA10 infected BALB/C mice. Consolidation of lungs parenchyma (C) denoted pathological state and Alveolar space (AS) denoted healthy state.

**Fig S2 Cap independent translation of SARS-CoV-2 and localization of La protein** (A) Mock and infected HEK-293T-ACE2 cells were harvested at 24h and fractionated for nucleus and cytoplasm. Representative western blot depicted distribution of La protein in nucleus and cytoplasm upon mock and infection. GAPDH was probed as cytoplasmic marker, and histone was probed as nucleus marker. (B) Representative graph depicted densitometric analysis of La protein distribution for nucleus and cytoplasm upon mock and infection conditions. (C) Represented graph depicted normalized luciferase value (Rluc / Fluc) using 5’UTR bicistronic constructs of SARS-CoV-2, Coxsackie virus B3 (CVB3) and EMCV. Null bicistronic construct was used as control. (D) *in vitro* transcribed capped and uncapped 5’UTR genomic and 5’UTR sub-genomic RNA of SARS-CoV-2 were transfected in HEK-293T-ACE2 cells and 12h post transfection cells were harvested for luciferase RNA and protein level estimation. Represented graph represented Normalized Rluc value, where R luc value was normalized by Rluc RNA level.

**Fig S3 Characterization of cleavage of La protein** (A) La protein was probed by NBP1 SSB antibody. Representative image depicted detection of La cleavage upon virus infection. (B) HEK-293T-ACE2 cells were infected with delta and omicron variant of SARS-CoV-2 at MOI 1 and cells were harvested 24h post infection. Representative image depicted detection of La cleavage upon delta and omicron infection.

**Fig S4 Molecular simulation of cleaved La and 5’UTR SARS-CoV-2 RNA** (A) IDR ensemble structure generated for N-terminal truncated La protein using IDR conformer generator and stitched with C-terminal truncated La protein and energy minimized structure generated for full length La. Representative Image depicted more stable IDR_4 full length structure. (B, C) Representative structural models of the final bound complexes of full-length La–5’UTR RNA and cleaved La–5’UTR RNA after MD simulations. (D, E) Backbone RMSD and RMSF analyses of RNA-bound complexes showing higher global deviation and altered residue flexibility in the cleaved La–RNA complex relative to the full-length complex. (F, G) Free energy landscape (FEL) projections illustrating a well-defined, thermodynamically favorable energy basin for the full-length La–RNA complex, whereas the cleaved construct samples broader conformational space.

**Table 1 Binding analytics of different dimers of La**

Table represented Gibbs free energy, dissociation constant and other parameters calculated for full length dimer, cleaved-full dimer, and cleaved-cleaved dimer.

**Table 2 M/GBSA and MM/PBSA binding energies of full length and cleaved La**

Table represented detailed energy parameters calculated for simulated full length and cleaved la protein in its bound form with 5’UTR of SARS-CoV-2.

