## Supplementary Figures for "Viral Protease Nsp5 Hijacks La Autoantigen to Orchestrate the Translation to Replication Transition in SARS-CoV-2"

A.

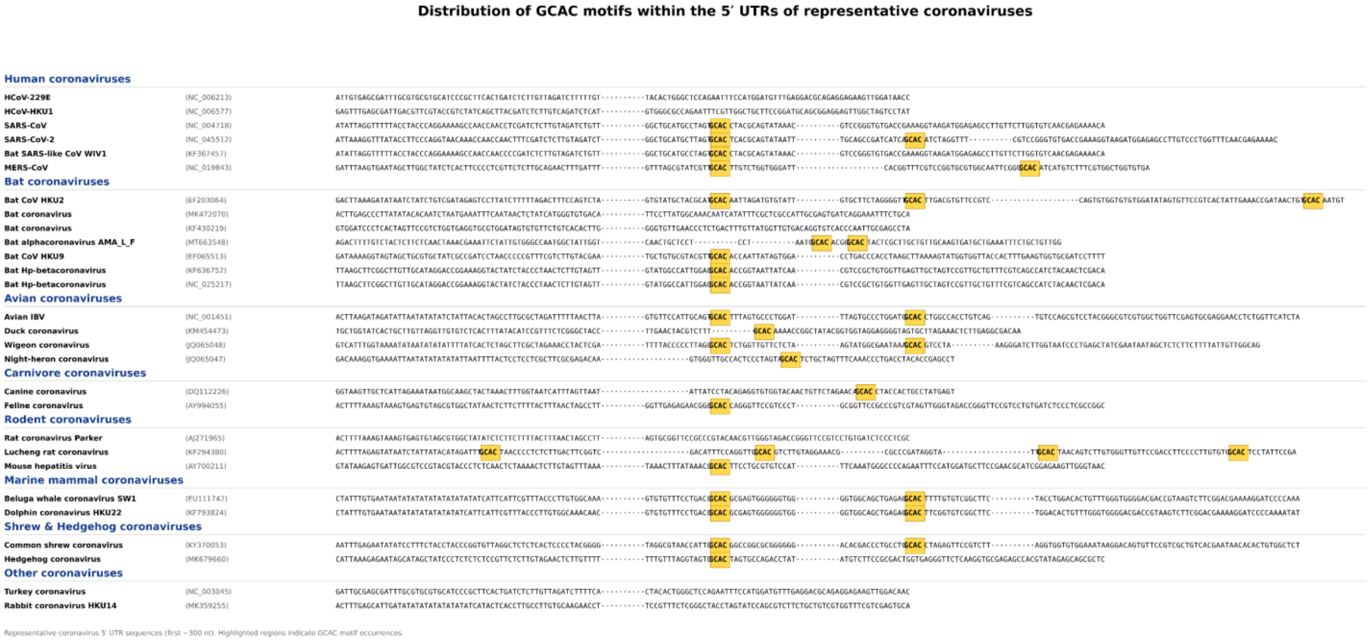


B.

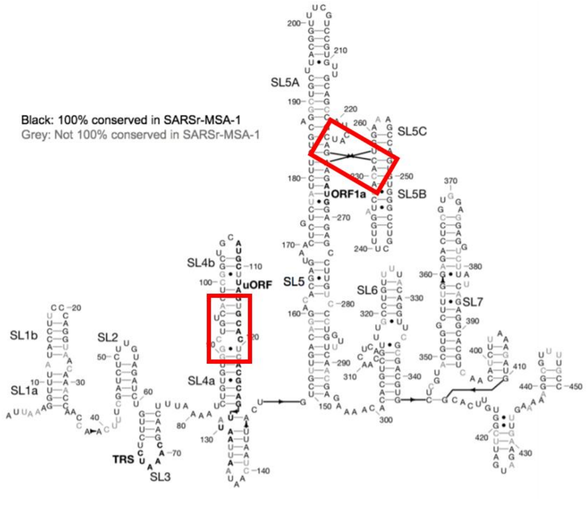


C.

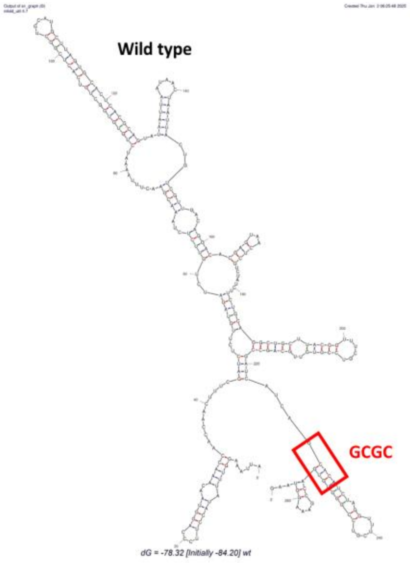


D.

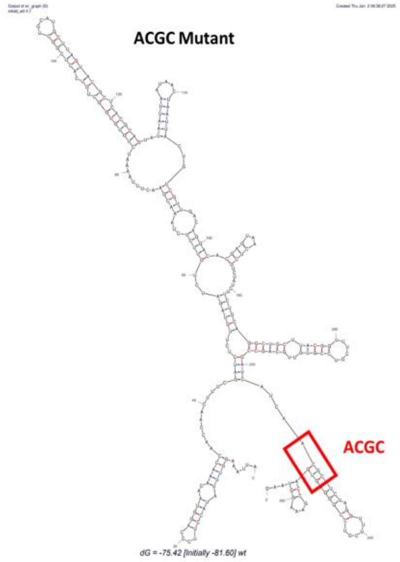


E.

```
Mus       MAENGDNEKMTALEAKICHQIEYYFGDFNLPRDKFLKEQIKLDEGWVPLETMIKFNRLNR  60
Homo      MAENGDNEKMAALEAKICHQIEYYFGDFNLPRDKFLKEQIKLDEGWVPLEIMIKFNRLNR  60
          **********:*************************************** *********

Mus       LTTDFNVIVQALSKSKAKLMEVSADKTKIRRSPSRPLPEVTDEYKNDVKNRSVYIKGFPT  120
Homo      LTTDFNVIVEALSKSKAELMEISEDKTKIRRSPSKPLPEVTDEYKNDVKNRSVYIKGFPT  120
          *********:*******:***:* **********:*************************

Mus       DATLDDIKEWLDDKGQILNIQMRRTLHKTFKGSIFAVFDSIQSAKKFVEIPGQKYKDTNL  180
Homo      DATLDDIKEWLEDKGQVLNIQMRRTLHKAFKGSIFVVFDSIESAKKFVETPGQKYKETDL  180
          ***********:****:***********:******.*****:******* ******:*:*

Mus       LILFKEDYFAKKNEERKQSKVEAKLKAKQEHEGRHKP-GSTETRALEGKMGCLLKFSGDL  239
Homo      LILFKDDYFAKKNEERKQNKVEAKLRAKQEQEAKQKLEEDAEMKSLEEKIGCLLKFSGDL  240
          *****:************.******:****:*.::*   .:* ::** *:**********

Mus       DDQTCREDLHFLFSNHGEIKWVDFAGGAKEGIILFKEKAKEALEKARNANNGNLLLRNKK  299
Homo      DDQTCREDLHILFSNHGEIKWIDFVRGAKEGIILFKEKAKEALGKAKDANNGNLQLRNKE  300
          **********:**********:**. ***************** **::****** ****:

Mus       VTWKVLEGHAEKEALKKITDDQQESLNKWKSKGGHAGGRFKGSHVFTAARRFKGKGKGNR  359
Homo      VTWEVLEGEVEKEALKKIIEDQQESLNKWKSKG----------------RRFKGKGKGNK  344
          ***:****..******** :*************                **********:

Mus       PGYAGAPKGRGQFHGRRTRFDDDD--------RRRGPMKRGRDGRDREEPASKHKKRENG  411
Homo      AAQPGSGKGKVQFQGKKTKFASDDEHDEHDENGATGPVKRAREETDKEEPASKQQKTENG  404
           .  *: **: **:*::*:* .**          **:**.*:  *:******::* ***

Mus       ARDK  415
Homo      AGDQ  408
          * *:
```

F.

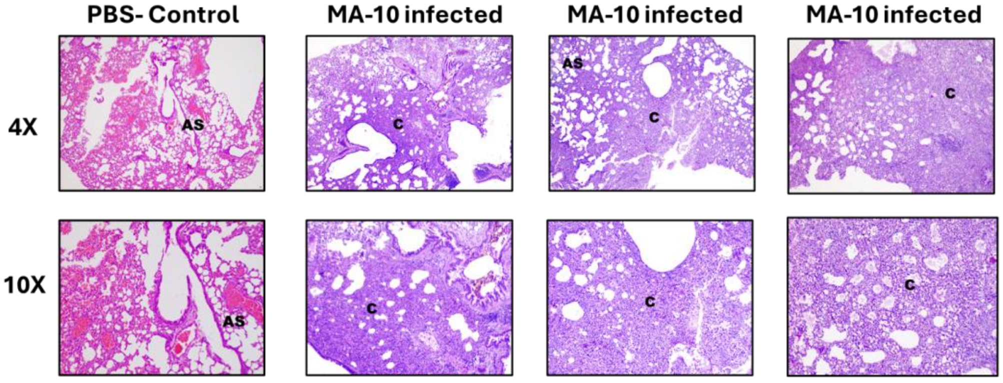


C-consolidation of lungs parenchyma (pathological state)
AS- Alveolar Space

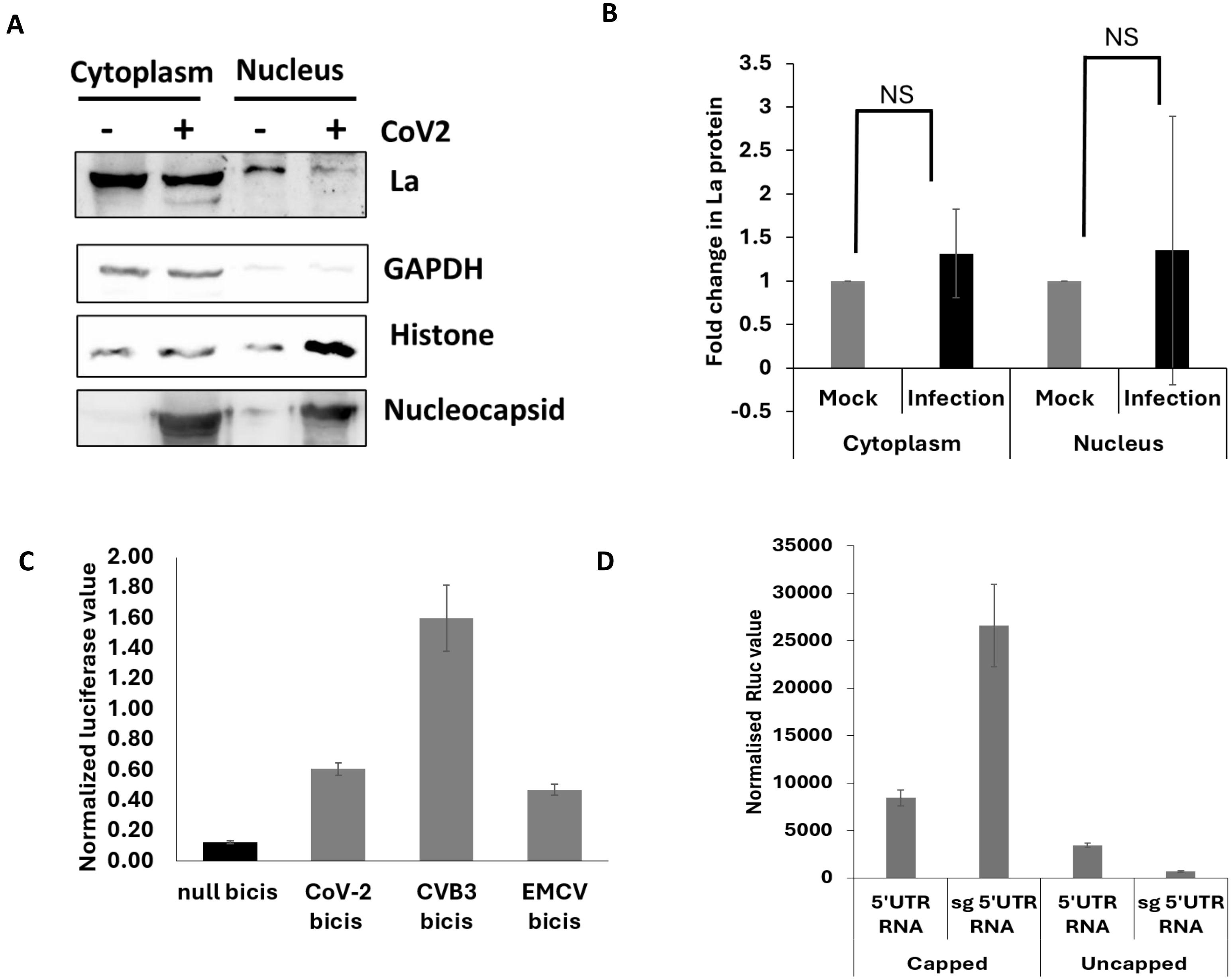
A
Cytoplasm
Nucleus
- + - + CoV2
La
GAPDH
Histone
Nucleocapsid
B
NS
NS
Fold change in La protein
3.5
3
2.5
2
1.5
1
0.5
0
-0.5
Mock
Infection
Mock
Infection
Cytoplasm
Nucleus
C
Normalized luciferase value
2.00
1.80
1.60
1.40
1.20
1.00
0.80
0.60
0.40
0.20
0.00
null bicis
CoV-2 bicis
CVB3 bicis
EMCV bicis
D
Normalised Rluc value
35000
30000
25000
20000
15000
10000
5000
0
5'UTR RNA
sg 5'UTR RNA
5'UTR RNA
sg 5'UTR RNA
Capped
Uncapped

**A**

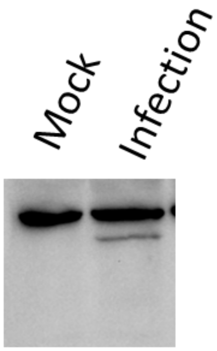


* NBP1 SSB antibody
(Novus biologicals)

**B**

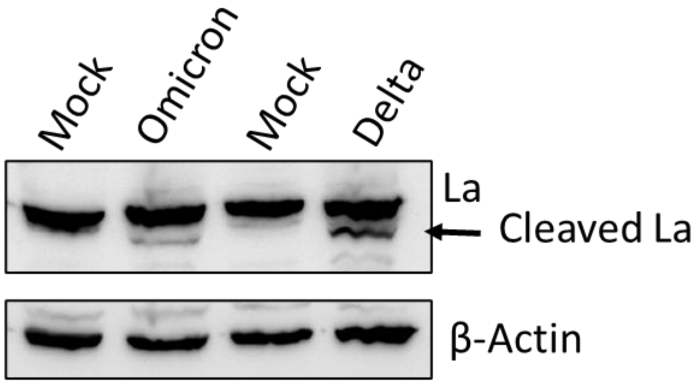

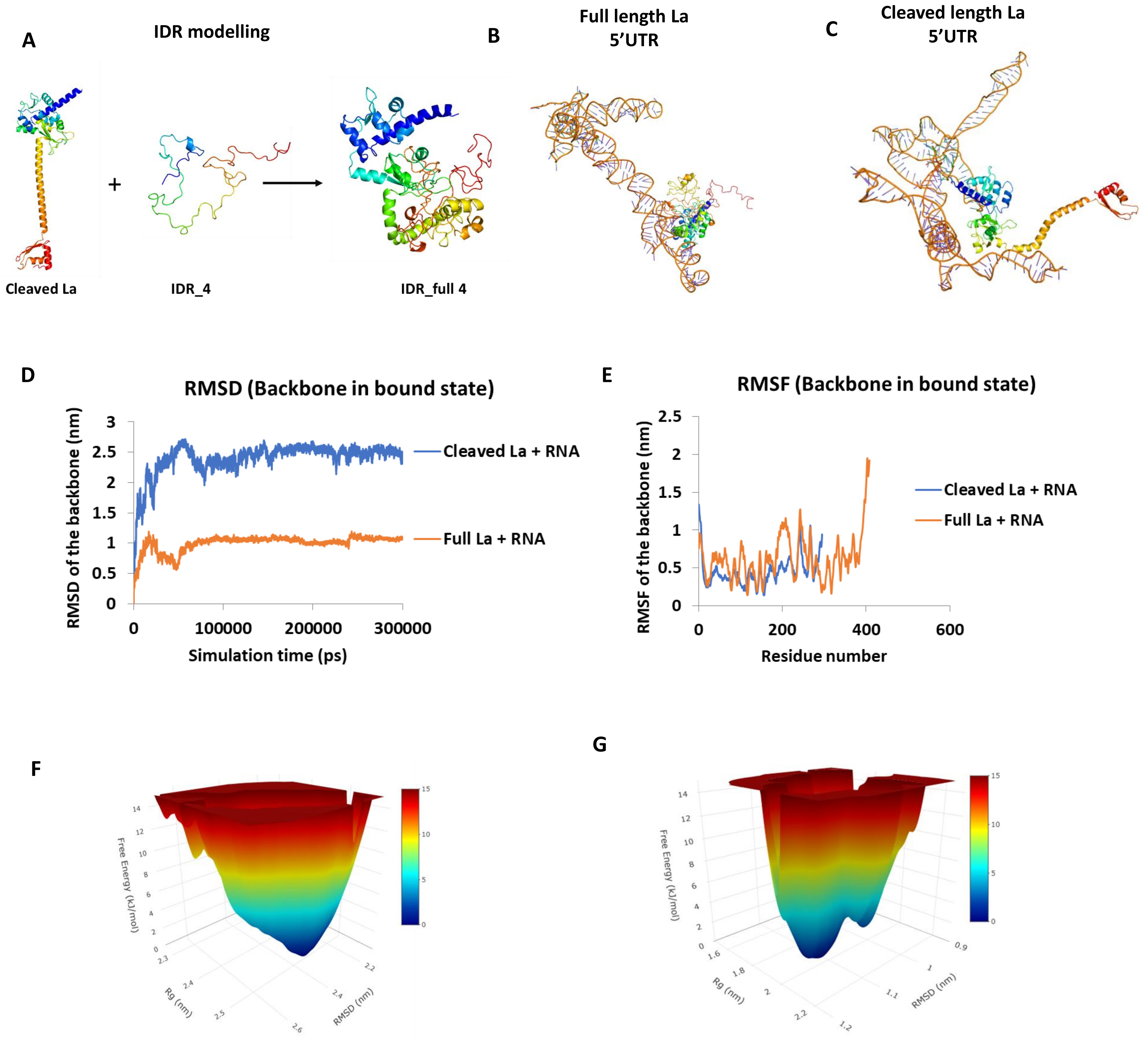
A
IDR modelling
Cleaved La
IDR_4
IDR_full 4
B
Full length La
5'UTR
C
Cleaved length La
5'UTR
D
RMSD (Backbone in bound state)
RMSD of the backbone (nm)
Simulation time (ps)
Cleaved La + RNA
Full La + RNA
E
RMSF (Backbone in bound state)
RMSF of the backbone (nm)
Residue number
Cleaved La + RNA
Full La + RNA
F
Free Energy (kJ/mol)
Rg (nm)
RMSD (nm)
G
Free Energy (kJ/mol)
Rg (nm)
RMSD (nm)

### TABLE 1

**Table – Binding analytics of different dimers of La**

| Protein-protein complex | ΔG (kcal mol-1) | Kd (M) | ICs charged-charged | ICs charged-polar | ICs charged-apolar | ICs polar-polar | ICs polar-apolar | ICs apolar-apolar |
|---|---|---|---|---|---|---|---|---|
| full_dimer | -14.5 | 2.50E-11 | 28 | 21 | 40 | 2 | 20 | 27 |
| cleaved_full_dimer | -7.7 | 2.30E-06 | 8 | 15 | 18 | 5 | 11 | 6 |
| cleaved_dimer | -6.1 | 3.30E-05 | 4 | 7 | 4 | 7 | 14 | 2 |

### TABLE 2

| | *MM-GBSA* | | *MM-PBSA* | |
|---|---|---|---|---|
| Energy Term | Full-Length La | Cleaved La | Full-Length La | Cleaved La |
| ΔVDWAALS | −196.65 ± 1.01 | −69.09 ± 0.52 | −196.65 ± 1.01 | −69.09 ± 0.52 |
| ΔEEL | −14504.26 ± 21.90 | +202.65 ± 39.86 | −14504.26 ± 21.90 | +202.65 ± 39.86 |
| ΔEGB | +14585.50 ± 21.40 | −164.42 ± 38.46 | +14496.48 ± 21.39 | −155.01 ± 38.31 |
| ΔESURF | −27.29 ± 0.10 | −9.57 ± 0.07 | −25.41 ± 0.08 | −8.33 ± 0.05 |
| ΔGGAS | −14700.91 ± 21.70 | +133.56 ± 39.73 | −14700.91 ± 21.70 | +133.56 ± 39.73 |
| ΔGSOLV | +14558.21 ± 21.40 | −174.00 ± 38.44 | +14471.07 ± 21.41 | −163.34 ± 38.32 |
| ΔG Total | −142.70 ± 1.08 | −40.44 ± 1.79 | **−229.84 ± 2.36** | **−29.78 ± 1.85** |

**Full-Length La (1–408) vs Cleaved La (1–295)**
All values in kcal/mol (mean ± SEM)
